# More than just axons: A positive relationship between an intracellular isotropic diffusion signal and pubertal development in white matter regions in a massive adolescent cohort

**DOI:** 10.1101/2022.11.23.517748

**Authors:** Benjamin T. Newman, James T. Patrie, T. Jason Druzgal

**Affiliations:** Department of Radiology and Medical Imaging, School of Medicine, University of Virginia; Department of Public Health Sciences, School of Medicine, University of Virginia

**Keywords:** Diffusion, Microstructure, MRI, development, puberty

## Abstract

Puberty is a key event in adolescent development that involves significant, hormone-driven changes to many aspects of physiology including the brain. Understanding how the brain responds during this time period is important for evaluating neuronal developments that affect mental health throughout adolescence and the adult lifespan. This study examines diffusion MRI scans from the cross-sectional ABCD Study baseline cohort, a large multi-site study containing thousands of participants, to describe the relationship between pubertal development and brain microstructure. Using advanced, 3-tissue constrained spherical deconvolution methods, this study is able to describe multiple tissue compartments beyond only white matter (WM) axonal qualities. After controlling for age, sex, brain volume, subject handedness, scanning site, and sibling relationships, we observe a positive relationship between an isotropic, intracellular diffusion signal fraction and pubertal development across a majority of regions of interest (ROIs) in the WM skeleton. We also observe regional effects from an intracellular anisotropic signal fraction compartment and extracellular isotropic free water-like compartment in several ROIs. This work suggests that changes during pubertal development elicit a complex response from brain tissue that cannot be completely described by traditional methods focusing only on WM axonal properties. This work brings in vivo human neuroimaging studies more into line with work performed on animal models, which describe an interaction between increased myelination, neurogenesis, angiogenesis, and glial cell proliferation in response to pubertal hormones.

## 1. Introduction

In adolescent development, puberty is a critical period that marks the hormonally driven transition into reproductive maturity^1^. Extensive physical, behavioral, and neurological changes occur rapidly and can have lifelong consequences for health and well-being. The developing brain undergoes substantial neuroanatomical reorganization at both global and cellular scales^2^. This critical period can leave the brain particularly vulnerable, and puberty marks the peak emergence of a majority of neuropsychiatric disorders^3,4^. Understanding the relationship between the cells of the brain and pubertal maturation is an important component of adolescent health and well-being throughout the remainder of the lifespan^2,5^. Previous neuroimaging studies have largely focused on axonal changes, however the white matter regions of the brain contain numerous glial cell varieties and extracellular space that contributes to a diverse diffusion profile^6^. In this study we present evidence for a widespread positive relationship between pubertal stage and an isotropic, intracellular, gray matter-like diffusion signal within the white matter skeleton. This signal may be indicative of the beginning of pubertal neuronal reorganization, with implications for understanding the neurobiology of maturing brain tissue.

Evidence from studies in animal models, where the levels of pubertal hormones can be directly manipulated, has demonstrated the effect of these hormones on the organization of brain tissue^7^. Pre-pubertal gonadectomy was shown to reduce the number of new, bromodeoxyuridine-positive cells in the brains of male and female rats, localized in a sex-dependent manner to regions that show post-pubertal sex-dependent differences in volume^8^. In female songbirds, axonal organization in song motor pathways have been reversibly manipulated via a testosterone injecting implant^9^. Pubertal hormones can influence the proliferation and survival of neurons mediated by local production of vascular endothelial growth factor and brain-derived neurotrophic factor^10^. It is unknown to what degree these processes are analogous to human development, as *in vivo* human studies must largely rely on neuroimaging techniques for insight.

Structural neuroimaging studies have consistently described a pattern of brain changes following the onset of puberty: an increase in the global volume of white matter (WM) and a decrease in the volume of gray matter (GM) in the cortex, sub-cortical nuclei, and at the WM-GM boundary^11–13^. It has long been thought that ‘thinning’ of cortical GM results from pruning unnecessary or inefficient synapses^14^. However, treating brain tissue as homogenous, binary, compartments to be examined individually may have obscured the process underlying the observed effect. The microstructural mechanism for this decrease in GM and increase in WM has been suggested to rely on increased myelination into the cortical GM areas which then appear to shift the location of the GM/WM boundary on T_1_-weighted images^15^. This possibility highlights the need for neuroimaging methods that can examine within-tissue cellular characteristics.

Diffusion MRI (dMRI) has provided more detailed measurements of *in vivo* brain development and allows for characterization of cellular microstructure within WM areas. Early dMRI studies using diffusion tensor imaging (DTI) found an increase in fractional anisotropy (FA), a measurement of axon coherence and myelination, and a decrease in mean diffusivity (MD) and radial diffusivity, indicative of reduced cellular permeability, as adolescents matured^11,16–18^. DTI is limited however, by a lack of specificity and from interference from heterogenous tissue composition or from multiple WM orientations (the ‘crossing fibers’ problem) in a single voxel^19,20^. By collapsing the diffusion signal into a single tensor, standard DTI metrics may be obscuring subtle but important cellular and sub-cellular changes in tissue composition.

More advanced dMRI analysis techniques – such as multi-compartment models and constrained spherical deconvolution methods – have improved on DTI by characterizing signals from multiple tissue types and/or multiple WM fiber orientations. Two studies have examined the early adolescent period using multi-compartment neurite orientation dispersion and density imaging (NODDI) which can provide more detailed measurements of brain microstructure than DTI^21^. Both studies found a widespread positive association with age in neurite density index, a model of intra-neurite space thought to describe myelination and axonal growth, but no relationship between age and orientation dispersion index, a model of inter-neurite space^22,23^.

Fixel-based analysis (FBA) is a promising dMRI model that builds off of spherical harmonic representations of WM that can overcome the pitfalls of DTI by being both sensitive and specific to WM fiber bundles without interference from isotropic signal from cell bodies/somas or from extra-cellular CSF^24,25^. A pair of studies have used FBA to examine a cross-sectional^22^ and longitudinal sample^26^ of developing subjects from the Children’s Attention Project study^27^. These studies explored the relationship between pubertal development, as measured by the pubertal development scale^28,29^, and WM measurements derived from FBA in ROIs defined by the Johns Hopkins University International-International Consortium Based Mapping (JHU-ICBM) atlas^30–32^. The cross-sectional study only found a significant relationship between fiber density and pubertal development in the splenium of the corpus callosum, while the longitudinal study found a broader range of significant areas throughout the WM skeleton.

In this study, we aim to move away from the WM axon focused metrics toward a more holistic measurement of brain microstructure accounting for multiple cellular environments encountered in the brain: axonal, anisotropic, WM-like diffusion; intracellular, isotropic, GM-like diffusion; and extracellular, isotropic, CSF-like diffusion^33,34^. Each of these compartments allow for the evaluation of different and distinct cellular environments within the same voxel. This study examines the relationship between pubertal development and cellular microstructure at the beginning of adolescence within ‘WM’ areas but measuring the whole microstructural environment, not exclusively axonal characteristics, as previous literature has done.

## 2. Materials and Methods

### 2.1 Participants

This study utilized baseline data obtained from the Adolescent Brain Cognitive Development (ABCD) Study, a publicly available neuroimaging and demographic study examining child development and factors leading to substance abuse. The ABCD study is the largest adolescent neuroimaging dataset ever acquired in the United States with an enrollment of 11,874 subjects at baseline. From this we selected a subset of 7,219 subjects enrolled at sites equipped with Siemens manufacturer scanners for processing and analysis. Following an automated quality control process including removing subjects without completed diffusion scans, excessive motion, or other imaging artifacts, we were able to successfully analyze 5,245 subjects in native, scanner space. Following template construction successful registration and warp into the template space was assessed by visual inspection aided by a semi-automated process and 4752 subjects remained for final analysis. The ABCD Study includes an oversampling of families with twins and the final cohort included 1070 twins, 19 triplets (drawn from 7 families of triplets, but in two families one subject did not pass quality control) and 563 non-twin siblings^35^.

### 2.2 Imaging data

Unprocessed dMRI images were obtained from the ABCD study and were acquired with a multiband accelerated sequence that had an isotropic voxel size 1.7 × 1.7 × 1.7 mm^3^ with TE = 88 ms and TR = 4100 ms. Using a multi-shell protocol 7 images were acquired at b= 0, 6 directions were acquired at b=500 s/mm^2^, 15 directions were acquired at both b=1000 s/mm^2^ and at b=2000 s/mm^2^, and 60 directions were acquired at b=3000 s/mm^2 36^. Two bidirectional field maps with reverse phase encoding were obtained at b=0 with identical isotropic voxel sizes, TE, and TR for use in distortion correction. Only images acquired using the Siemens Prisma 3T platform were analyzed to avoid manufacturer and sequence differences, including field gradient strength, TE, and TR, that have previously been demonstrated to affect outcome 3 Tissue, Constrained Spherical Deconvolution model (3T-CSD) signal fraction results^34^.

Total brain volume was obtained from the ABCD study which released processed data as part of the baseline data release. Study organizers obtained this volumetric data from T1 weighted images with an isotropic voxel size 1.0 × 1.0 × 1.0 mm^3^ with TE = 2.88 and TR = 2500 with a flip angle of 8 degrees and an field of view of 256 × 256 mm^36^. Volumetric data was calculated using an automated processing pipeline in Freesurfer and released as part of the ABCD data release 2.0.1. Specific details of ABCD image processing have been published previously^37^.

### 2.3 dMRI Image Preprocessing and Analysis

Image preprocessing was performed consistent with prior protocols that have been shown to result in consistent and reliable signal fraction measurements^34^. All dMRI images were corrected for thermal noise using the “dwidenoise” command implemented in MRtrix3^38^. Gibbs rings were then removed with the “dwidegibbs” MRtrix3 function^39^. The FSL package (“topup” and “eddy”) was subsequently applied to correct for susceptibility-induced (EPI) distortions, eddy currents, and subject motion, including the Gaussian replacement of outliers^40–43^. Finally the preprocessed images were upsampled using MRtrix3 to alter the resolution to 1.3 × 1.3 × 1.3 mm^3^ isotropic voxel size, in order to bring their spatial resolution closer to the high-resolution acquisition of images in the Human Connectome Project which serves as a benchmark for high quality diffusion data^44,45^. The b=0 and b=3000 s/mm^2^ shells were then extracted to form a single-shell image set suitable for single-shell, 3 tissue constrained spherical deconvolution (SS3T-CSD). This step was performed because prior investigations have suggested SS3T-CSD is superior at differentiating between brain regions compared to multi-shell multi-tissue CSD at b=3000 s/mm^2^ ^34^.

Brain masks were obtained for all subjects by performing a recursive application of the Brain Extraction Tool^46^. Response functions from each of the three tissue types were estimated from a randomly selected subset of nearly 500 subjects and averaged to produce a single set of tissue response functions^47^. The response functions were selected via an unsupervised method described by Dhollander et al., (2016) where the WM tissue response function was selected from an FA thresholded mask, the CSF tissue response function was selected in voxels with the highest signal decay metric between the averaged b-0 and b=3000 s/mm^2^ shells, and the GM tissue response function was selected from voxels closest to the median voxel-wise signal decay metric, after a conservative GM mask was constructed. SS3T-CSD was then performed using the average response functions with MRtrix3Tissue, a fork of MRtrix3, to estimate: 1) an anisotropic WM-like compartment (represented by a complete WM fiber orientation distribution (FOD), i.e. all fit signal from WM fibers), 2) an isotropic, intracellular GM-like compartment (the signal that is fit by neither WM or CSF response functions, and more closely fit by an isotropic but less b-value dependent signal response function), and 3) an isotropic, extracellular CSF-like compartment^34,48^. SS3T-CSD is functionally a specialized optimizer that iteratively performs a linear least squares fit of the response functions similarly to multi-shell multi-tissue CSD (MSMT-CSD). Initially, the isotropic CSF and GM response functions are fit to the diffusion signal, with WM as a prior constraint. This calculation yields an underestimate of CSF from the CSD fit. The next step fits anisotropic WM and isotropic GM response functions with the previously calculated CSF as a constraint, also yielding an underestimate of WM from the CSD fit. Over continuous iterations, this method attempts to assign signal repeatedly to either WM or CSF compartments from the GM compartment, repeatedly performing a cleaner separation as signal is assigned and remaining signal is run through the algorithm again. An example of this general pipeline, including preprocessing steps, is available at https://3tissue.github.io/doc/single-subject.html. Each subject’s three tissue compartments were then normalized to sum to 1 on a voxel-wise basis, resulting in the final three-tissue signal fraction maps^33,34^. Summing the spherical harmonic coefficients on a rotational-invariant voxel-wise basis provides the added benefit of harmonizing inter-scanner and inter-subject signal intensity differences while preserving between-subject biological variation^49^.

A cohort specific template was constructed from a random selection of 50 subjects’ WM-FODs (Fig. 1a) using symmetric diffeomorphic registration of the FOD themselves and implemented in MRtrix3 with the “population_template” function^50^. Each subject was then individually registered to the cohort template using an affine, followed by a nonlinear registration guided by the WM FODs themselves in an unbiased manner. The resulting warp was used to move each of the subjects’ three-tissue signal fraction maps into template space. Quality control was performed on the transformed signal fraction maps via visual confirmation of a semi-automated method that flagged images with abnormal values.

**Figure 1:**
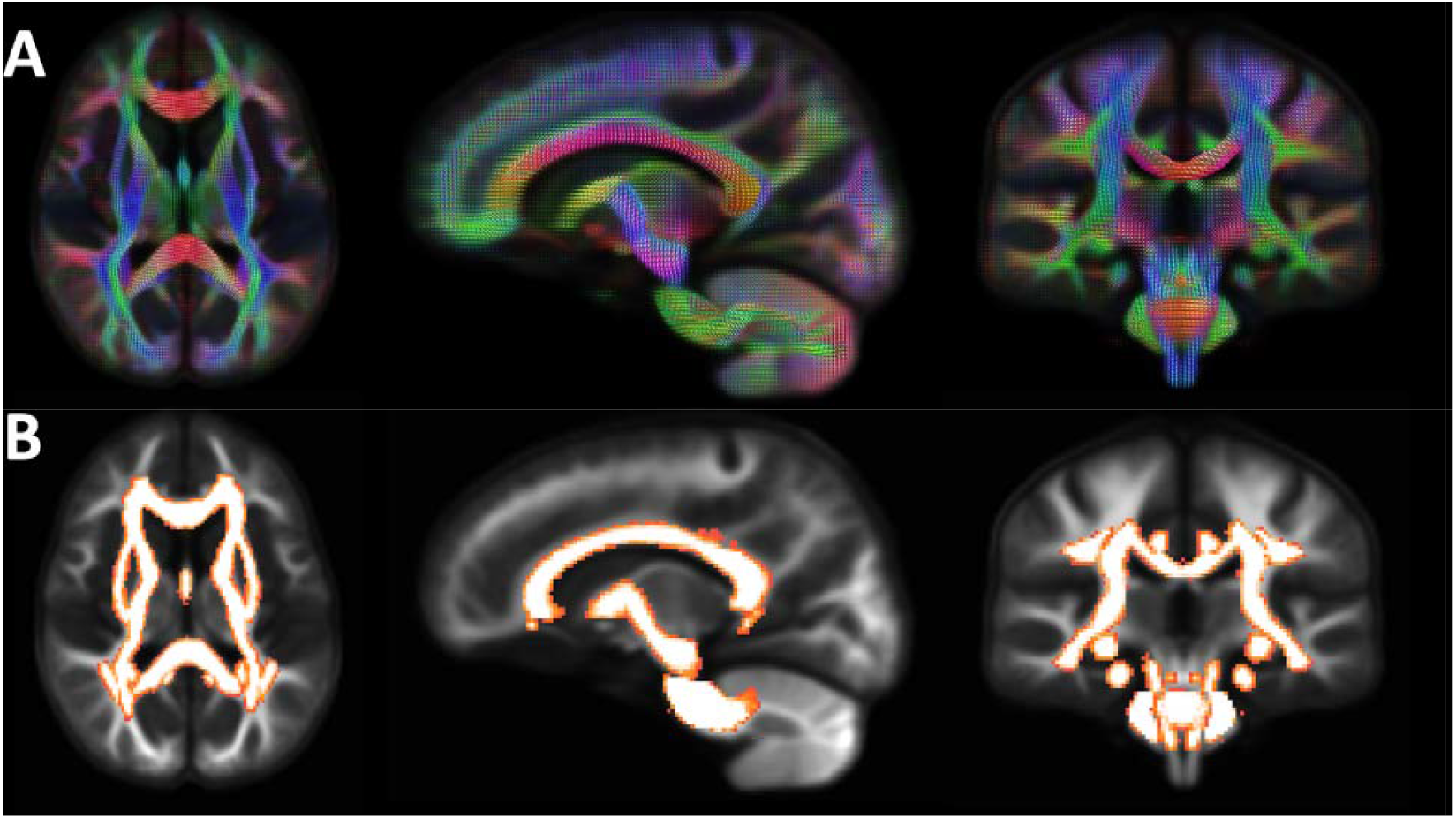
Graphic displaying the WM FODs of the cohort specific group template constructed from the average of 50 randomly selected ABCD subjects. The top row (A) displays the FODs themselves while the bottom row (B) shows the positioning of the 48 ROIs in the JHU-ICBM atlas after being warped into the template.

The cohort template was then registered to stereotaxic space with a similar FOD-based diffeomorphic registration procedure to a b-value matched version of the NTU-DSI-122 template^51^. The resulting warp from stereotaxis space was used to move the 48 regions of interest (ROIs) included in the JHU-ICBM DTI-81 white matter atlas (available as part of FSL, hereafter referred to as the JHU WM atlas; Fig. 1b) into the cohort specific template space^30–32,52^. The average value of each signal fraction map within each of the 48 ROIs was calculated using the “mrstats” function from MRtrix3.

### 2.4 Pubertal Development Scale

Measures of pubertal development were assessed via parental completion of the Pubertal Development Scale (PDS) questionnaire during the baseline subject visit^28,53^. The PDS measures physical manifestations of pubertal development on a scale from 1 (no development) to 4 (development seems complete) across 3 measures common to both sexes and 2 measures each for adrenarche and gonadarche features. A total PDS score (PDSS) was calculated for each individual by summing the combined common, adrenarche, and gonadarche features of the questionnaire^29^. PDSS has been demonstrated to significantly correlate with saliva testosterone, DHEA, and estradiol levels^29^, was recorded for nearly every ABCD subject, and allows for more simplified direct comparison between sexes compared to saliva or serum hormone levels.

### 2.5 Statistical Analyses

#### Data summarization

Categorical data are summarized by frequencies (n) and percentages (%). Continuous scaled data are summarized by the mean, standard deviation, and range of the empirical distribution.

#### Patient demographic and patient characteristic analyses

A one-sample binomial exact test was performed to compare gender cross-sectional frequencies. A two-sample t-test was performed to compare the age distributions of the female and male cross-sectional samples. A two-sample t-test was also performed to compare the subject-specific average puberty score between female and male children.

#### Tissue signal fraction regression analyses

A mixed effect model regression was performed to predict the mean ROI tissue signal fraction (i.e. extracellular isotropic CSF-like; or intracellular isotropic GM-like; or intracellular anisotropic WM-like) in a given JHU atlas ROI as a function of the sum of the child’s PDS scores (PDSS-sum), the child’s age (years) and sex (female, male), the child’s handedness (left, right, and ambidextrous) and the child’s total brain volume (cm^3^). Each child’s family and sibling relationship was accounted for as a nested random effect in the model to ensure twins, triplets, and siblings were not biasing results following recommendations by Saragosa-Harris et al., (2022)^54^. Site effects were controlled for as an additional random effect. All of the model predictor variables and interactions that were selected a priori based on scientific merit, and the model is displayed below as Equation 1.

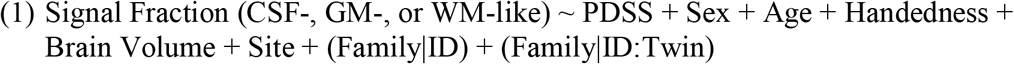

For each JHU atlas ROI, the concomitant variable adjusted association between tissue signal fraction (extracellular isotropic CSF-like, or intracellular isotropic GM-like, or intracellular anisotropic WM-like) and PDSS-sum was quantified by the regression slope coefficient estimate associated with PDSS-sum. For each JHU atlas ROI, a null hypothesis test was performed to test the null hypothesis that the slope of the association between the tissue signal fraction and PDSS-sum is equal to 0, versus the alternative that the slope of the association between the tissue signal fraction and PDSS-sum is not equal to 0. The complete set of p-values from the 48 different JHU atlas ROI were then subjected to the Benjamini and Hochberg false discovery procedure to identify those ROI, in which the ANOVA type II F-test p-value of the null hypothesis test was less than the Benjamini and Hochberg 0.05 false discovery threshold. This correction was performed for each 48 ROIs separately as the signal fraction results from each ROI are interdependent (i.e. they sum to one in each voxel) and are separate models to aid interpretation.

## 3. Results

### 3.1 Participant Demographics

There were significantly more males than females in the cross-sectional sample (2496 males (52.5%) versus 2256 females (47.5%), p<0.001) and significantly more right-handed subjects (right = 3806 (80.2%) versus left = 628 (13.2) versus ambidextrous = 315, p<0.001 for all). The age range was the same for the female cohort and the male cohort (107.0 to 132 months), and there was no significant difference between the average age of females (119.5 months ± 7.4 SD) and the average age of males (119.9 months ± 7.5 SD) (p=0.086). As predicted by the early age range of participants, the distribution of the pubertal development scale score (PDSS) was not uniformly distributed across the range of PDSS values (i.e. 1 to 4). The mean of the distribution for average PDSS was 1.62 units (95% CI: [1.60, 1.63]). There was a highly significant relationship between average PDSS and age (slope = 0.118 PDSS/month; 95% CI: [0.096, 0.141], p<0001); which remained true for both females (slope = 0.212 PDSS/year; 95% CI: [0.181, 0.244], p<0.001) and males (slope = 0.049; 95% CI: [0.02, 0.078], p=0.001), but the slope for rate in change in average PDSS/month was greater for females than for males (p<0.001). Females also had a significantly higher average PDSS score compared to males (1.77; 95% CI: [1.76, 1.80] versus 1.47; 95% CI: [1.45,1.49], respectively, p<0.001) (Fig. 2).

**Figure 2:**
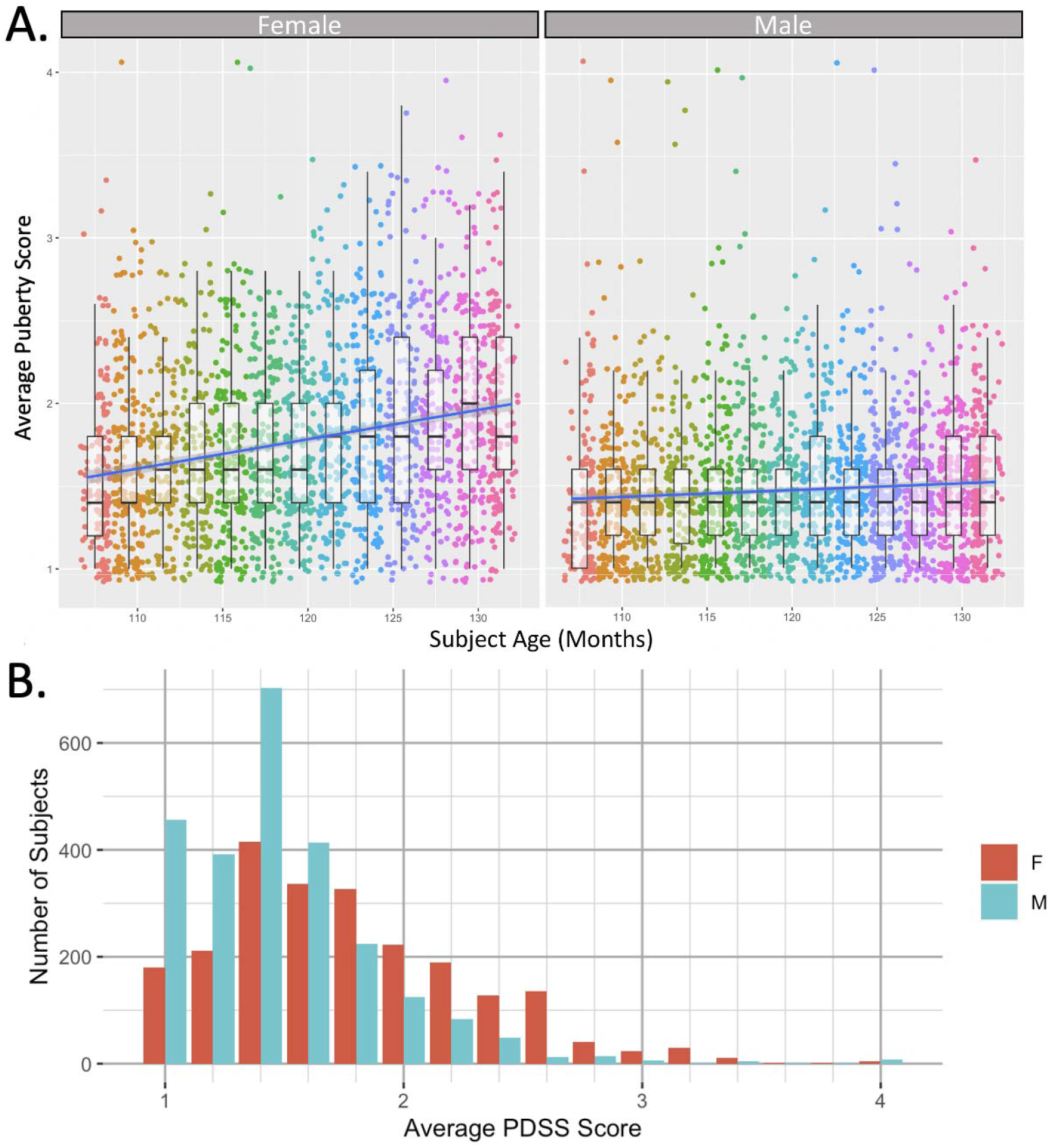
The relationship between subject age and average PDSS score with each boxplot and corresponding color representing a 2-month bin of participants (A). A linear model is displayed (shading is SE) to illustrate the relationship between age and average PDSS score. Both male and female members of the participant cohort tended to have a higher mean PDSS score as age increased, with this positive relationship being significantly more pronounced for female participants compared to males. A histogram (B) shows similar trends in the distribution of PDSS scores separated by sex.

### 3.2 Regression Analyses

#### Tissue signal fraction regression analyses

The ANOVA summaries for the mixed-effect regression models that were utilized to account for random nested effects from sibling relationships and random effects from scanner site predict tissue signal fraction (i.e. extracellular isotropic CSF-like; or intracellular isotropic GM-like or intracellular anisotropic WM-like) as a function of the sum of the child’s PDSS scores, the child’s age and sex, the child’s handedness, and child’s total brain volume. The results of these models are summarized in Table 1 displaying the number of JHU ROI that had a significant association with each of the predictor variables. Of note is the widespread reliability of PDSS-score, age, sex, and total brain volume as significant predictors across multiple tissue types, particularly of the intracellular isotropic signal fraction compartment (GM-like). Also of note is the almost complete lack of handedness as a predictor of intracellular signal fraction compartments.

**Table 1:** Number of JHU ROI (out of 48 total) where each associated variable remained significantly associated with tissue signal fraction after implementation of the Benjamini and Hochberg false discovery error rate procedure (p<0.05 false discovery rate). Age, followed by total brain volume, then sex, was significantly associated with each tissue signal fraction across a majority of ROIs. PDSS score was significantly associated with only GM-like tissue signal fraction in a majority of ROIs, while handedness was not significantly associated with GM-like tissue signal fraction in any ROI.

| <u>Tissue Signal Fraction</u> | <u>PDSS score # ROI (%)</u> | <u>Sex # ROI (%)</u> | <u>Age # ROI (%)</u> | <u>Total Volume # ROI (%)</u> | <u>Handedness # ROI (%)</u> |
| --- | --- | --- | --- | --- | --- |
| **CSF-like** | 19 (39.6) | 27 (56.3) | 43 (89.6) | 35 (72.9) | 8 (16.7) |
| **GM-like** | 40 (83.3) | 39 (81.3) | 44 (91.7) | 43 (89.6) | 0 (0.00) |
| **WM-like** | 9 (18.8) | 38 (79.2) | 45 (93.8) | 44 (91.7) | 3 (6.25) |

### 3.3 Concomitant variable adjusted associations between tissue signal fraction and PDSS-sum

Table S1 lists the regression model adjusted slopes, 95% confidence intervals, and p-values for predicting tissue signal fraction as a function of PDSS sum when the regression model input values for child age and sex, the child handedness and child total brain volume are held constant. Table S2 lists the individual p-values that have been summarized in Table 1 for each concomitant variable after the Benjamini and Hochberg false discovery rate procedure. The intracellular anisotropic WM-like, intracellular isotropic GM-like, and extracellular isotropic CSF-like tissue signal fraction related adjusted slopes, and 95% confidence intervals atlas are graphically displayed for the 48 different JHU atlas ROIs in Figure 3, Figure 4, and Figure 5, respectively.

**Figure 3:**
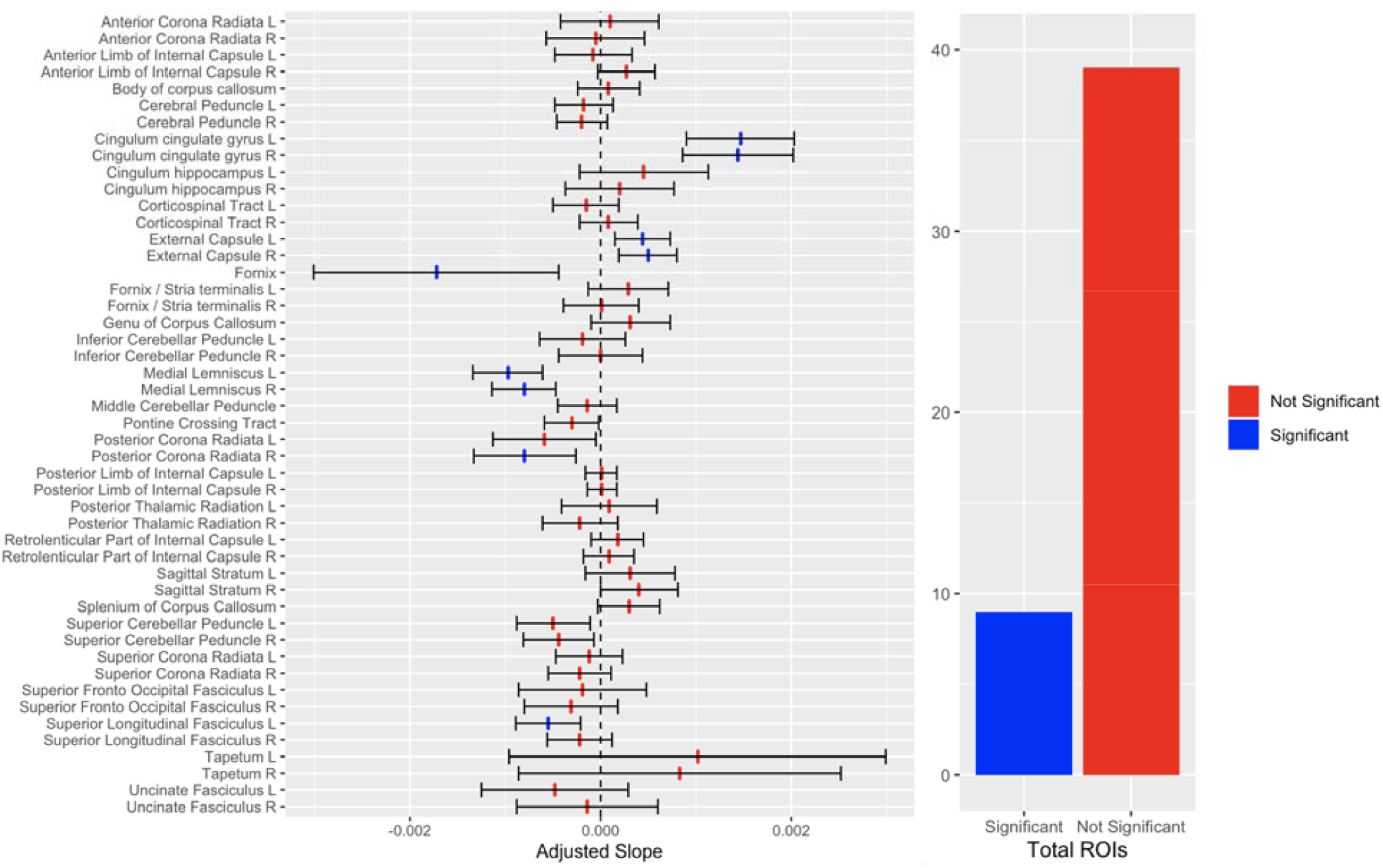
Chart displaying the adjusted slope with 95% confidence interval for the relationship between WM-like signal fraction and average PDSS score in each of the 48 JHU atlas ROIs. Blue bars indicate the model calculated a significant relationship (p<0.05) after Benjamini and Hochberg (1995) adjustment.

**Figure 4:**
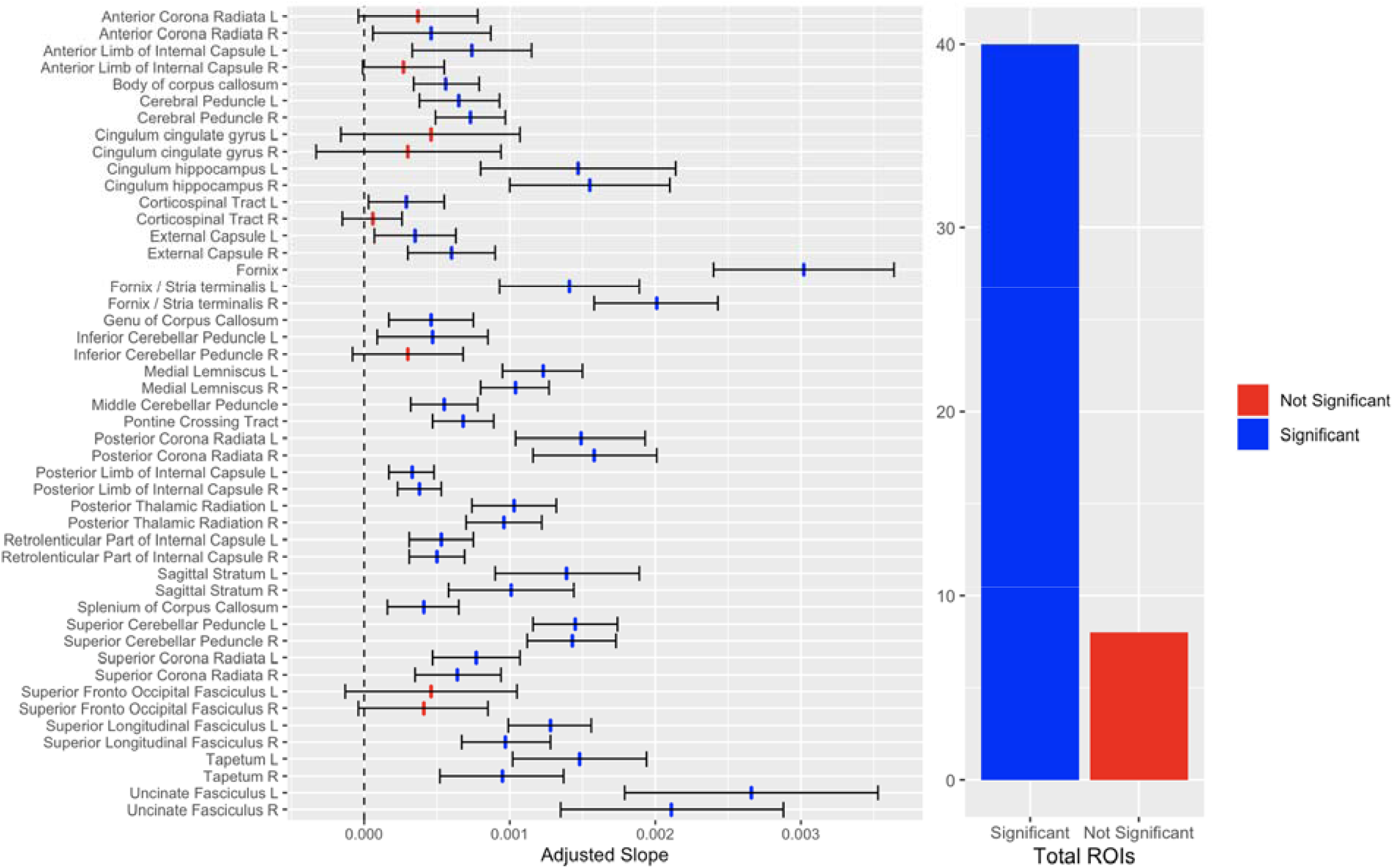
Chart displaying the adjusted slope with 95% confidence interval for the relationship between GM-like signal fraction and average PDSS score in each of the 48 JHU atlas ROIs. Blue bars indicate the model calculated a significant relationship (p<0.05) after Benjamini and Hochberg (1995) adjustment. The large majority of ROIs had a significant relationship between GM-like signal fraction and PDSS score, demonstrating a robust effect across the brain.

**Figure 5:**
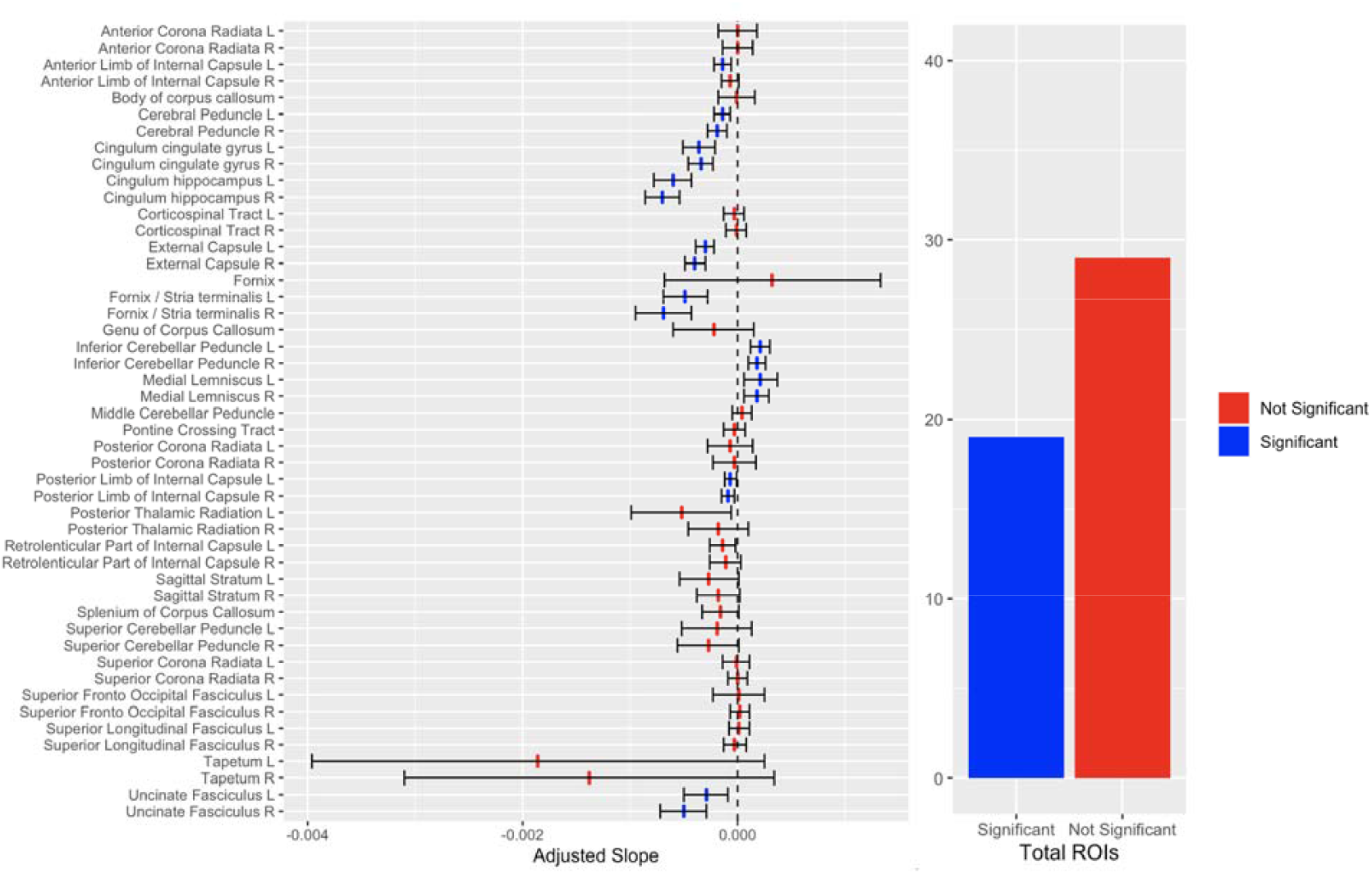
Chart displaying the adjusted slope with 95% confidence interval for the relationship between CSF-like signal fraction and average PDSS score in each of the 48 JHU atlas ROIs. Blue bars indicate the model calculated a significant relationship (p<0.05) after Benjamini and Hochberg (1995) adjustment. Just under half of ROIs examined displayed a significant relationship with PDSS score.

For each significant ROI from the intracellular isotropic GM-like tissue compartment the direction of the association is consistent, with intracellular isotropic GM-like signal fraction increasing with PDSS. For intracellular anisotropic WM-like and extracellular CSF-like signal fractions the direction of association was not consistent throughout the examined ROIs. The location of significant intracellular anisotropic WM-like, intracellular isotropic GM-like, and extracellular isotropic CSF-like tissue signal fraction adjusted model slopes are displayed in position on the cohort specific template in Figure 6, Figure 7, and Figure 8, respectively. Significant regions for both the intracellular tissue types tended to localize to the posterior portions of the brain, while significant extracellular regions located below the brainstem had a positive association with PDSS, and those above had a negative association with PDSS score.

**Figure 6:**
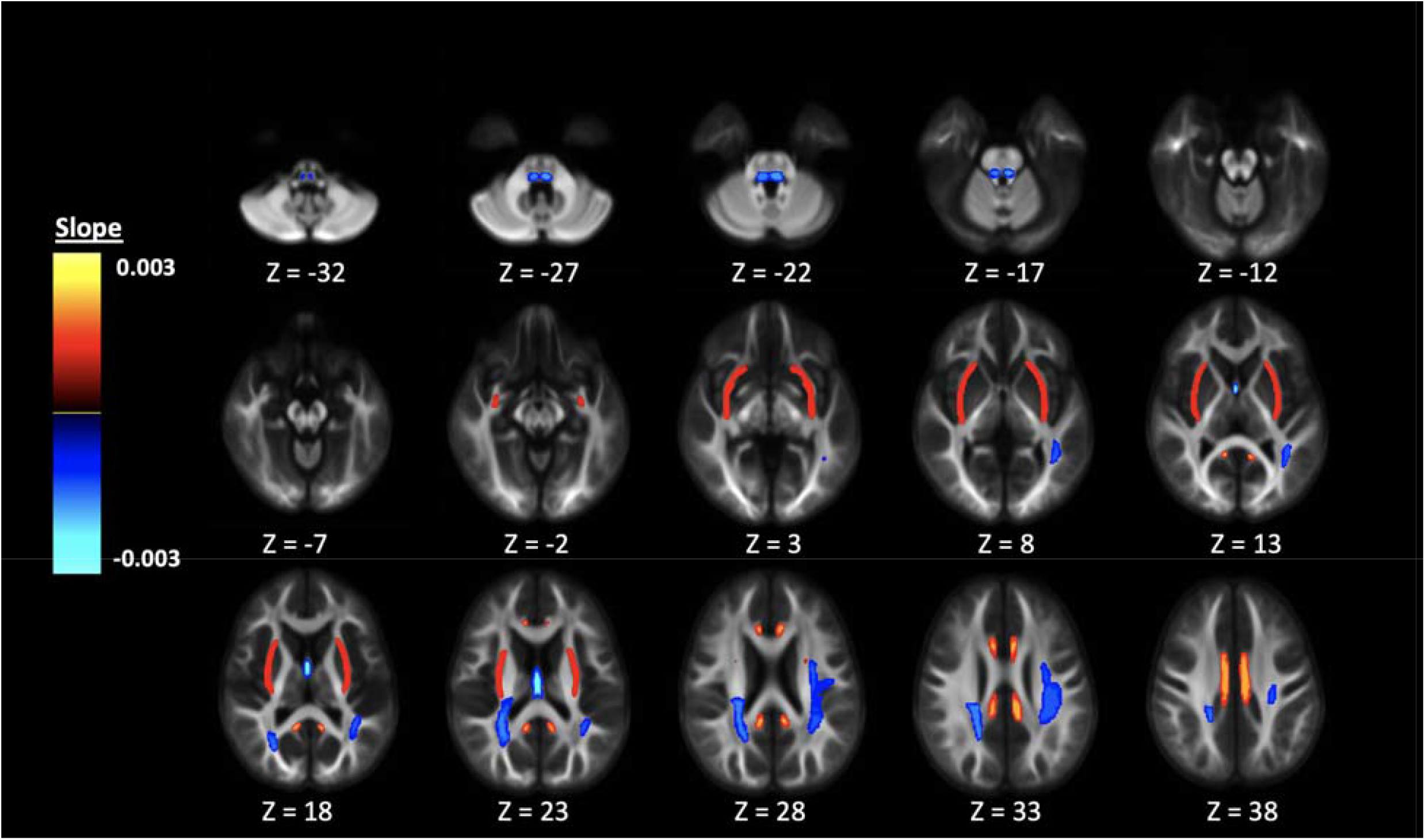
Display of significant adjusted anisotropic intracellular WM-like signal fraction model slopes from ROIs in the JHU WM atlas colored by slope and displayed on the cohort specific template. ROIs located in the posterior parts of the brain appear to be more strongly negatively associated with PDSS score than regions elsewhere.

**Figure 7:**
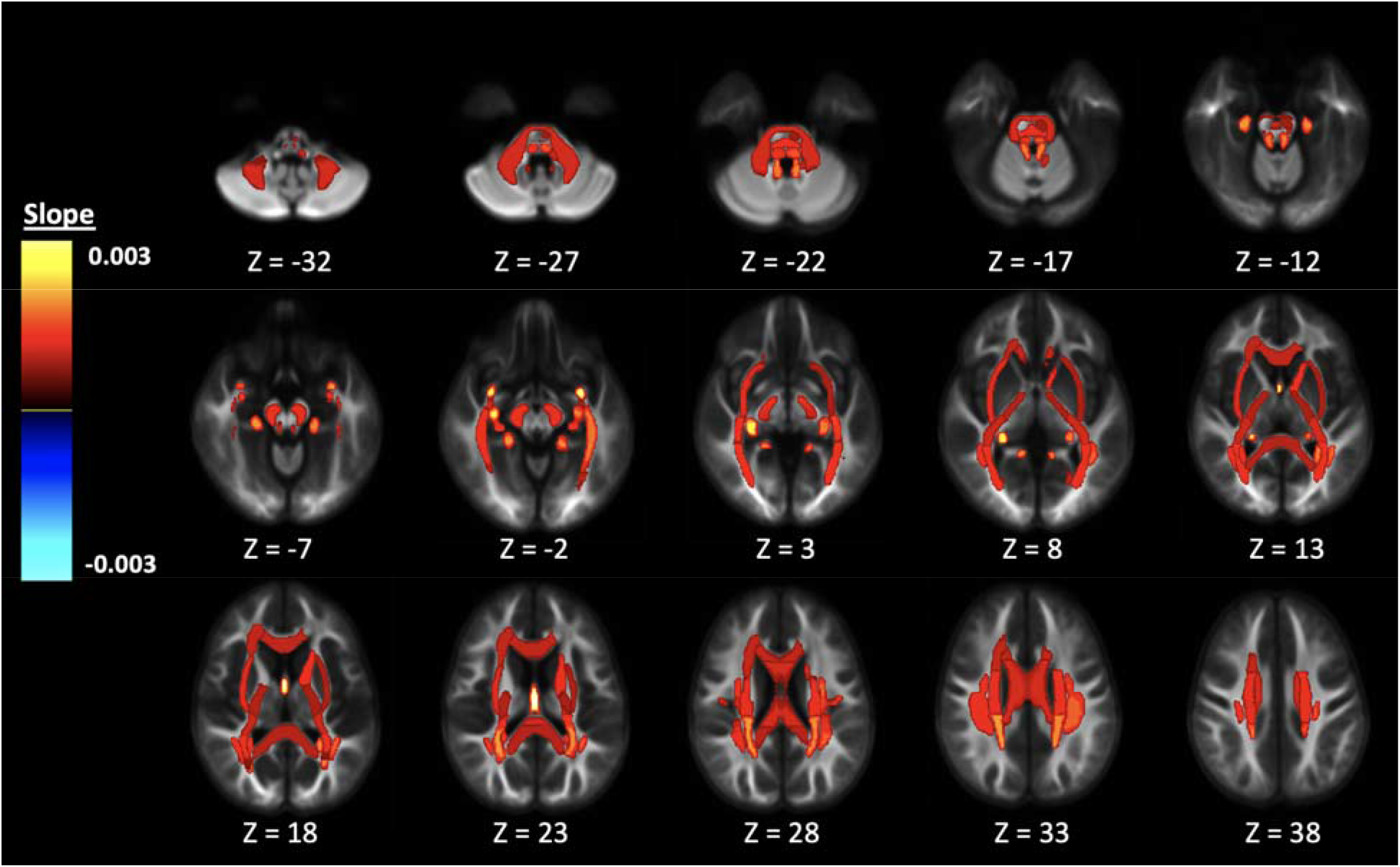
Display of significant adjusted isotropic intracellular GM-like signal fraction model slopes from ROIs in the JHU WM atlas colored by slope and displayed on the cohort specific template. ROIs located in the posterior parts of the brain appear to be more strongly positively associated with PDSS score than regions elsewhere.

**Figure 8:**
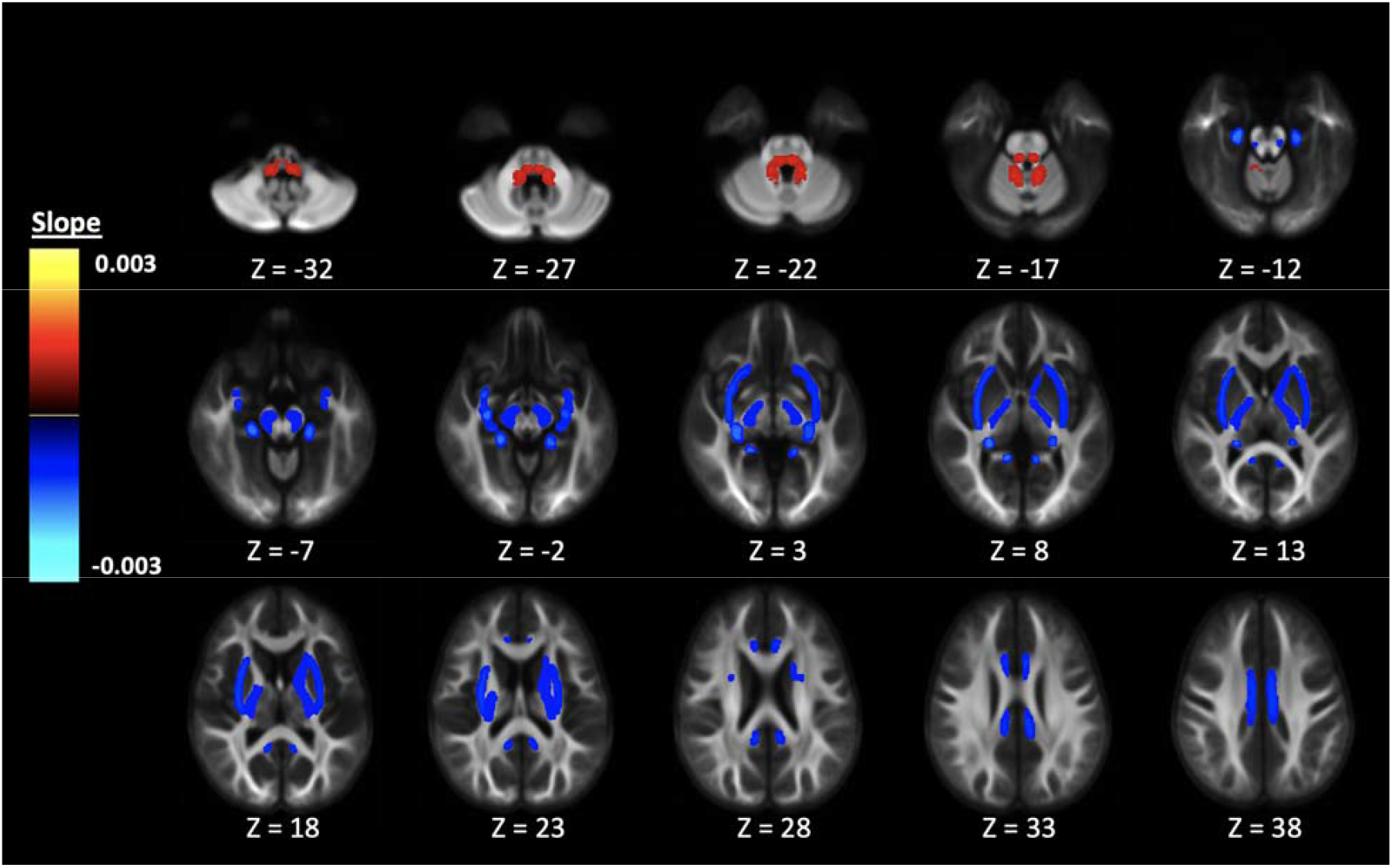
Display of significant adjusted isotropic extracellular CSF-like signal fraction model slopes from ROIs in the JHU WM atlas colored by slope and displayed on the cohort specific template. The CSF-like signal fraction was the only tissue compartment to have ROIs with significant adjusted model slopes both positive and negatively associated with PDSS score. Significant ROIs located below the brainstem had a positive association with PDSS, while those above had a negative association with PDSS score.

## 4. Discussion

In this cross-sectional diffusion MRI study of 4752 adolescents from the baseline collection of the ABCD study we have identified a relationship between physical manifestations of pubertal status and several measurements of brain tissue microstructure across a widespread number of WM brain regions. In ROIs for which the relationship was significant, 3T-CSD measures of intracellular anisotropic GM-like signal fraction were found to have a positive relationship with PDSS. The intracellular anisotropic WM-like and extracellular isotropic CSF-like signal fraction, in contrast, did not have a consistent direction across all significant ROIs.

There are several plausible mechanisms occurring at the cellular level of the brain that might underlie the effect observed in this study, but due to the nature of the signal fraction analysis each tissue compartment is flexible and nonspecific and precludes identifying a particular cellular type as the root cause of the observed changes. Possible sources for these cellular changes have been described in animal models. These vary from neurogenesis to the development of new astrocytes occurring during puberty, either of which could be a plausible reason for the observed GM-like signal fraction increase^7,8,55,56^. Neurogenesis and astrocytic growth would add to the intra-cellular space within a voxel without necessarily contributing immediately to the axonal volume and would thus appear as an increase in intracellular isotropic GM-like signal fraction at the expense of intracellular anisotropic WM-like signal fraction instead of extracellular isotropic CSF-like signal fraction (already relatively low in the predominantly WM JHU-ICBM atlas ROIs). Work performed using a songbird model observed angiogenesis and altered neuronal processes such as dendrite outgrowth, cell spacing, and an increased number of synaptic vesicles in response to increased levels of androgenic and estrogenic exposure^10^. Changes in cellular architecture due to pubertal hormones could underlie the changes observed in this study. Recent work has found relationships between CSD- and DTI-based measures of microstructure in song-specific brain areas and plasma testosterone levels, indicating that cellular changes due to pubertal hormones might be sufficiently large enough to be detected with dMRI metrics^9^.

It is also possible that our results largely reflect a critical period early in pubertal development due to the early age and pubertal status of our cross-sectional sample. Some studies have suggested that GM volume peaks around the age of our cohort. One DTI study found that for males in an early stage of pubertal development, as determined by hormone measures, there was a positive relationship between MD and age^57^. While they are not equivalent, increasing MD and increasing isotropic GM-like signal fraction could both occur from the same underlying cause in predominantly axonal areas, such as neurogenesis or increased cellularity. Given the ages of males in our study and the low average PDSS score for individuals in the group it is possible that this effect contributed to the observed results in this study.

This study also observed significant effects of age, sex, and total brain volume on the signal fraction measurements across the vast majority of ROIs. Several neuroimaging studies have not reported sex differences, or have reported a limited effect of sex, despite an earlier onset of puberty in girls. Genc et al.,^26^ for example, reported a significant effect of sex on fiber density in only one ROI. Similarly, a longitudinal study focusing on cortical thickness found changes associated with pubertal development and a significant effect of sex in only two ROIs^58^. The current study may have found a more widespread relationship across the brain, despite being cross-sectional in design, by including a great deal more participants (over 4,700, compared to 74 in Genc et al.,^22^ 130 in Genc et al.,^26^ and 126 in Herting et al.,^58^). Other large cohort studies, including those across longer longitudinal timeframes than in this study, have found similar differences between sexes. One study examining DTI metrics across a 5 year longitudinal sample of 8-28 year-olds’ found broad changes across a number of ROIs, suggesting that WM maturation is a gradual process that requires either large cohorts of subjects or long longitudinal data to fully detect^18^.

A recent study examined the same ABCD dataset used here in a longitudinal manner and applied a restriction spectrum imaging (RSI) model to examine changes in brain microstructure in relation to chronological age. The study found that a restricted compartment (thought to represent axonal signal) increased throughout the brain with age while a hindered compartment and a free water compartment had negative relationships with age^59^. While the present study primarily examined the effect of puberty, there was also widespread significant effects of age, perhaps suggesting that development is also driven through non-hormonal means. Additionally, while the RSI model shares many similarities with 3T-CSD, it is possible that the specifics of each model, such as the hindered compartment in RSI being modeled with a 4^th^ order spherical harmonic with physiological constraints while 3T-CSD uses a 0^th^ order spherical harmonic for the GM-like compartment, affect the physiological changes that contribute to each compartment. For example, narrower bodies such as neurites and glial processes may be better represented by the hindered compartment (which was developed on cortical animal tissue) while the GM-like compartment may represent isotropic bodies such as somas or cell bodies^60^. Histological validation of 3T-CSD is needed to make assumptions about underlying cellular processes and their effects on signal fractions.

The findings from the current study detailing an intracellular isotropic signal positively associated with puberty appear counterintuitive compared to prior studies examining the relationship between puberty and dMRI measures of brain microstructure, which have generally found that measures related to WM integrity (such as FA) increase while measures of permeability (such as MD) decrease^61^. This review includes studies using other similar CSD-based metrics such as apparent fiber density, a voxel-wise relative measure of intra-axonal volume^62^ or the similarly derived fiber cross section, both of which are termed ‘fixel-based metrics’ and provide specific quantitative information about the WM FOD^63^. Studies examining apparent fiber density have found significantly increased axonal fiber development related to the same PDSS measurement employed in our study^22,26^ and to testosterone^64^. Our results diverge from these studies in that the intracellular isotropic GM-like tissue compartment was positively associated with pubertal development throughout the brain while the intracellular anisotropic WM-like tissue compartment was much less commonly significantly associated in the same atlas and with the same pubertal measurements used previously^22,26^. It is important to note that the 3T-CSD technique measures relative levels of each tissue compartment and does not exclude the possibility WM fibers also mature in response to pubertal development. Possible physiological interpretations of this effect include changes in the number or activity of glial cells, such as oligodendrocytes responsible for increased myelin^65^, or apoptotic events that are thought to occur during neuronal pruning, which may contribute to an increased intracellular isotropic signal via the breakup of axons and their consumption by glial cells^66^.

Another study focusing on fixel-based metrics found a significant relationship between pubertal hormone levels and fiber density and fiber cross section across much of the posterior voxels in the JHU-ICBM atlas in a longitudinal cohort of children aged 8.5-10 years old^64^. While the pattern of significant voxels largely matches expectations from earlier structural studies^7^ and ROIs found to be significant in this study, there are several methodological reasons why this result may be similar to volumetric studies without shedding light on brain microstructure. The authors used a CSD method previously shown to systematically *overestimate* the contribution of WM fibers to the diffusion signal^48^ and the analyses did not control for longitudinal effects of change in brain volume. The combination of these two methodologic choices may be enough to cause some of the reported effects. These same methodologic factors could also plausibly explain the lack of sex effects, as well as the largely testosterone-based hormone relationship as testosterone has been reported as having a large effect on brain volume^67^.

Perhaps the most straightforward reconciliation between the fixel results and the present study is that fixel-based metrics are - at the voxel level - freely-varying descriptions of WM signal characteristics whereas 3T-CSD signal fractions are relative measurements of each tissue compartment’s contribution to total signal^24,34^. Fixel-based analysis discards the isotropic signal from CSF and GM in order to better characterize the WM signal while 3T-CSD examines how each tissue exists to some degree in every voxel. It is thus entirely possible that axonal microstructure matures in response to pubertal development, and that this process is accompanied by a local increase in cellularity with a GM-like intracellular diffusion profile. 3T-CSD is not specific enough to determine if this is caused by increased glial cell activity, neurogenesis, developing myelination, or a gross change in cellular architecture as axons reorganize, but it does suggest that a focus exclusively on WM is insufficient to understand the whole brain response to pubertal development.

While it is difficult to attribute the origin of the effects observed in this study to a specific cellular process, it is apparent that there is a broad and significant change within axonal regions of the brain in response to adolescent pubertal development. Using advanced dMRI measurements we have found a widespread positive relationship between an isotropic, intracellular GM-like signal fraction and pubertal development in a cross-sectional cohort. This finding has implications for the study of the cellular basis of human brain development, suggesting microstructure beyond axons or processes such as neurogenesis or phagocytosis contribute to adolescent brain WM development and are measurable by dMRI. This work also suggests that future adolescent neuroimaging studies should account for changes in non-axonal tissue compartments and pubertal development.

## 5. Conclusion

In this cross-sectional dMRI study of 4752 adolescents from the baseline collection of the ABCD study we have identified a relationship between physical manifestations of pubertal status and several measurements of brain tissue microstructure across a widespread number of primarily axonal brain regions. Our multicompartment 3T-CSD model agrees with evidence from animal models that cellular processes other than axonal signal are involved in white matter development and that future neuroimaging studies of WM in pubertal cohorts might benefit from a focus on cellular signal beyond axons.

## Supplementary Tables

**Table S1:**
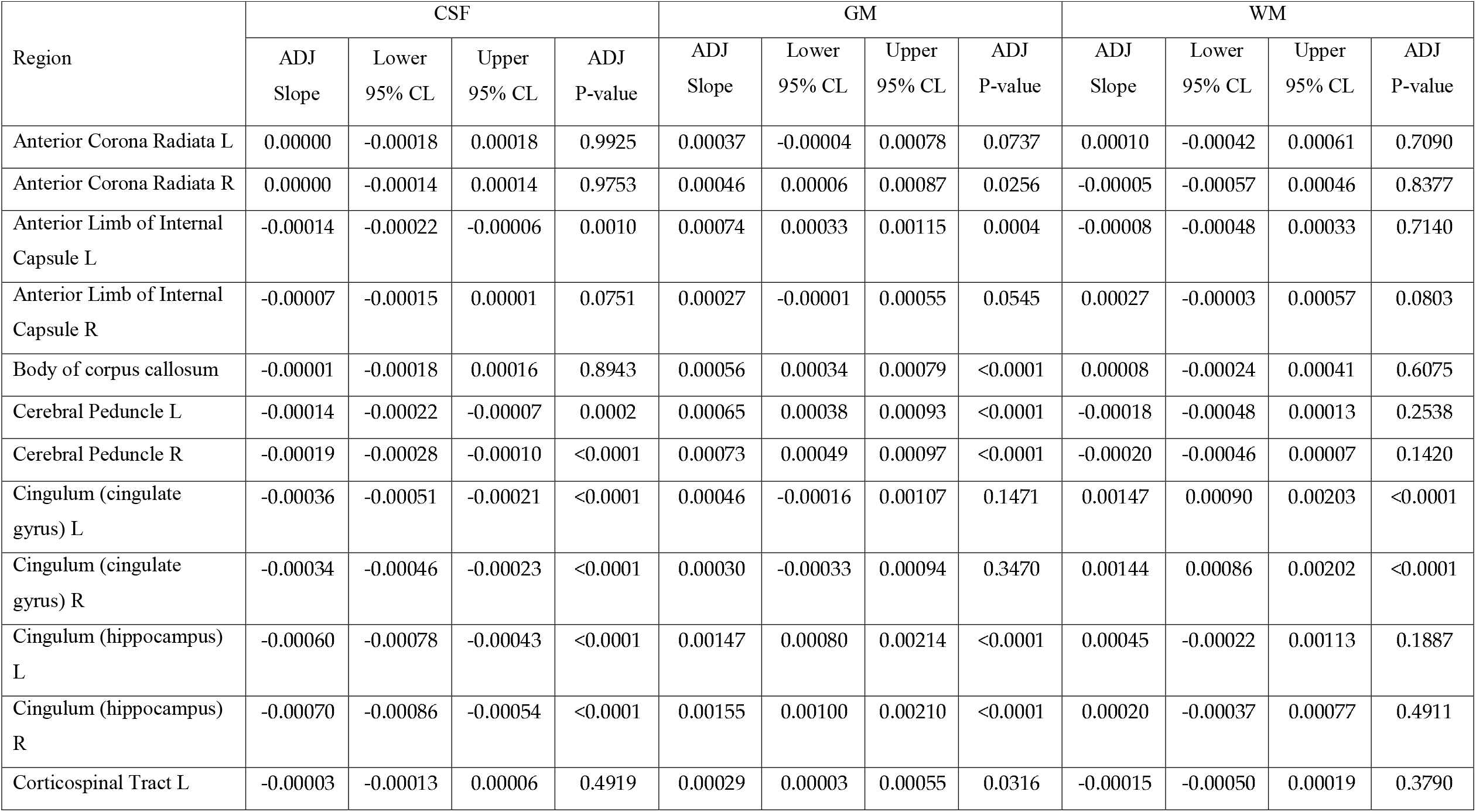
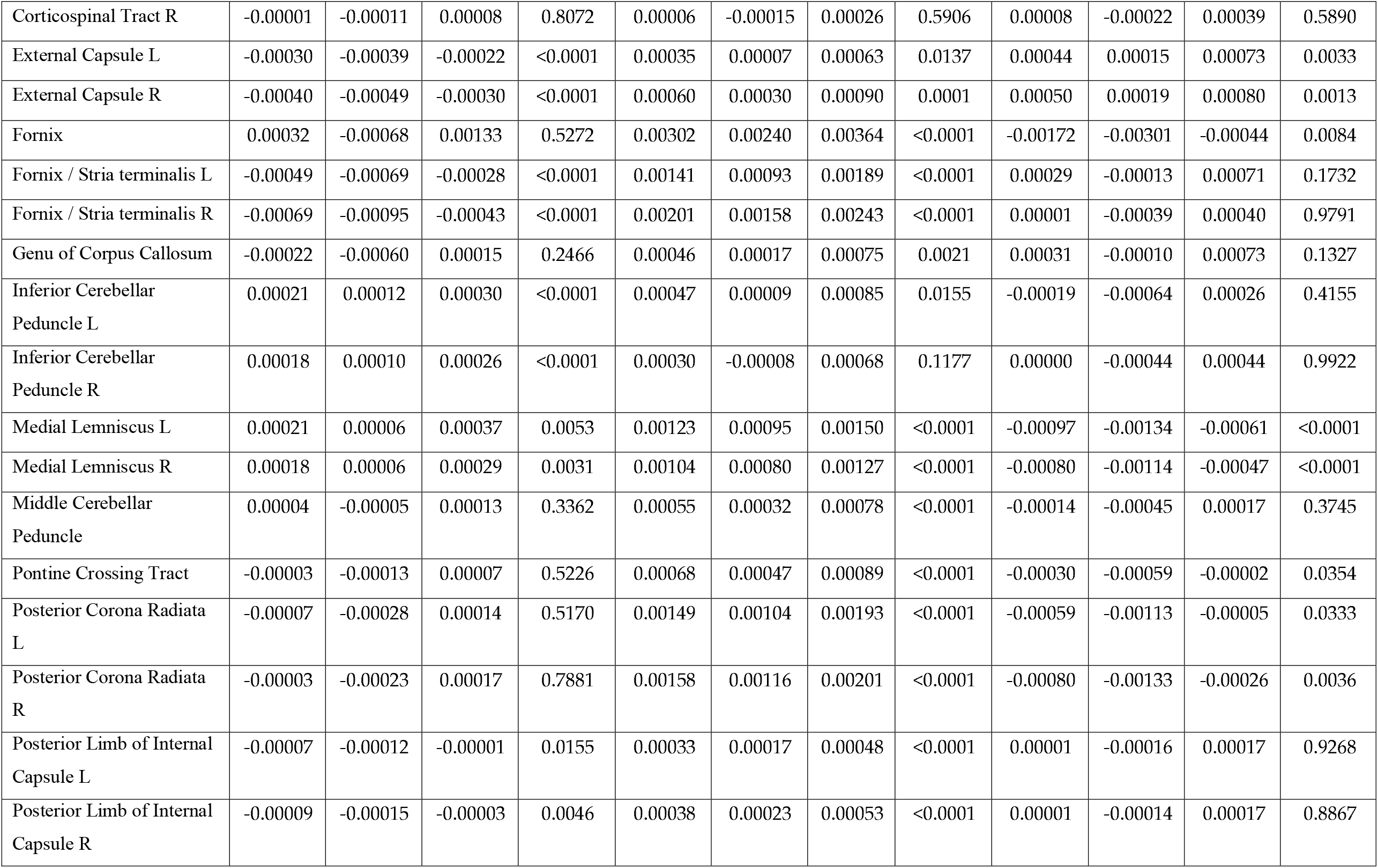
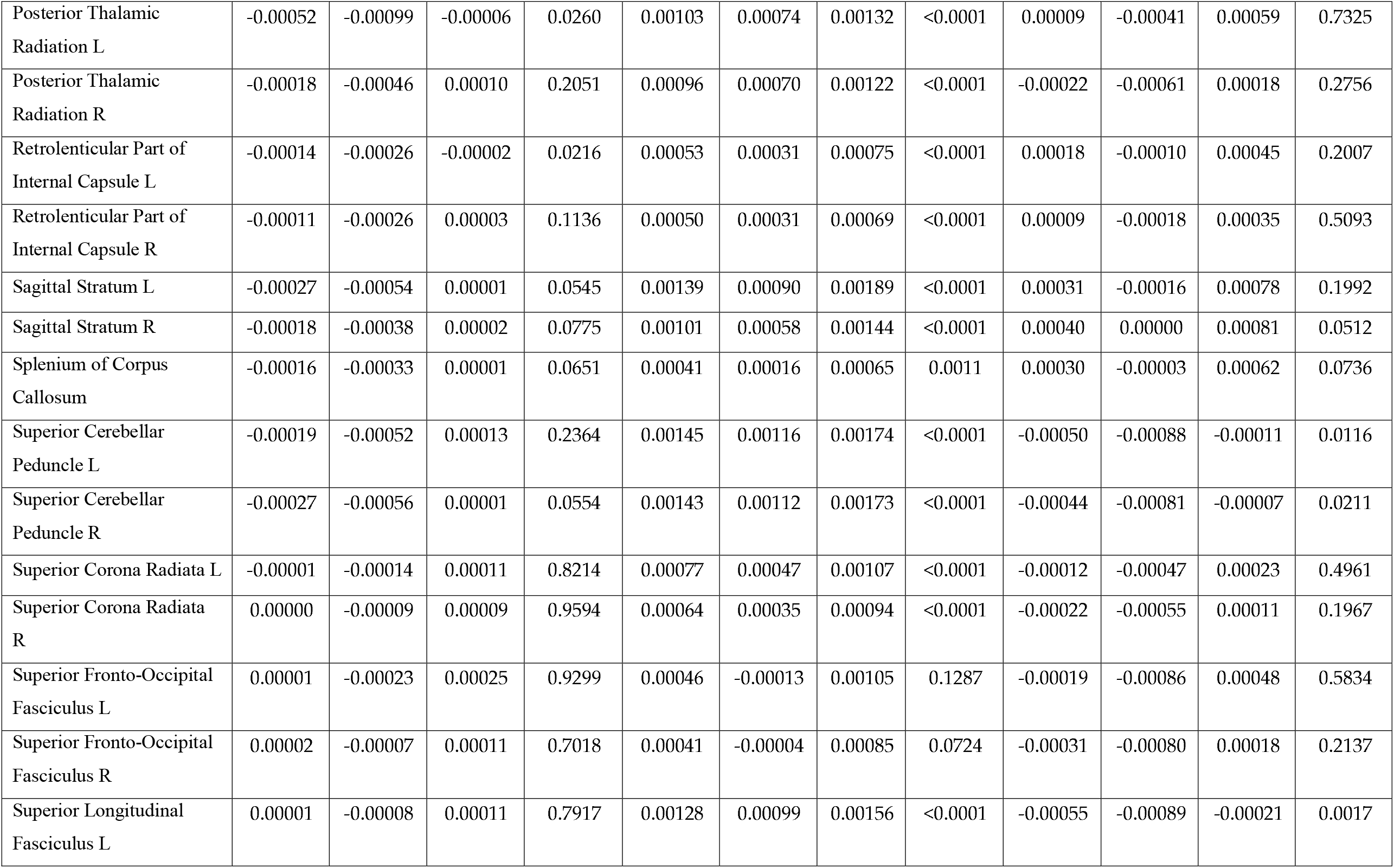
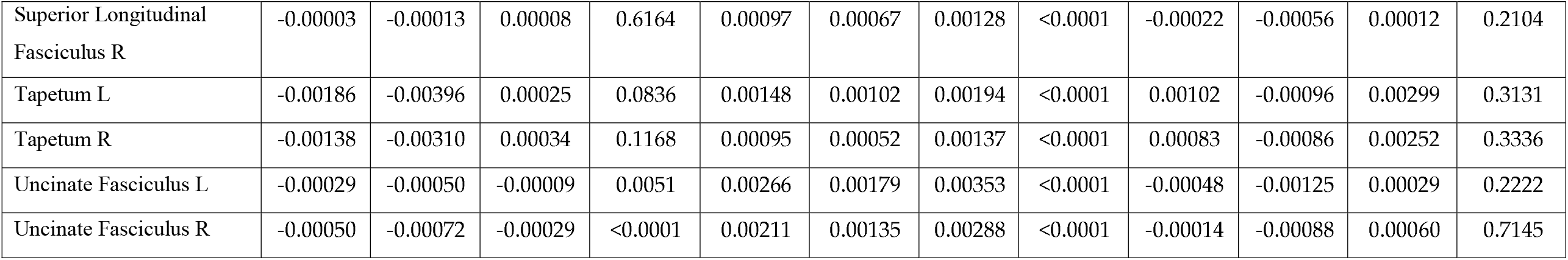
Adjusted slopes, 95% confidence levels, and p-values for the relationship between tissue signal fraction and PDSS sum in each of the JHU atlas ROIs.

| Region | CSF |  |  |  | GM |  |  |  | WM |  |  |  |
| --- | --- | --- | --- | --- | --- | --- | --- | --- | --- | --- | --- | --- |
|  | ADJ<br>Slope | Lower<br>95% CL | Upper<br>95% CL | ADJ<br>P-value | ADJ<br>Slope | Lower<br>95% CL | Upper<br>95% CL | ADJ<br>P-value | ADJ<br>Slope | Lower<br>95% CL | Upper<br>95% CL | ADJ<br>P-value |
| Anterior Corona Radiata L | 0.00000 | -0.00018 | 0.00018 | 0.9925 | 0.00037 | -0.00004 | 0.00078 | 0.0737 | 0.00010 | -0.00042 | 0.00061 | 0.7090 |
| Anterior Corona Radiata R | 0.00000 | -0.00014 | 0.00014 | 0.9753 | 0.00046 | 0.00006 | 0.00087 | 0.0256 | -0.00005 | -0.00057 | 0.00046 | 0.8377 |
| Anterior Limb of Internal Capsule L | -0.00014 | -0.00022 | -0.00006 | 0.0010 | 0.00074 | 0.00033 | 0.00115 | 0.0004 | -0.00008 | -0.00048 | 0.00033 | 0.7140 |
| Anterior Limb of Internal Capsule R | -0.00007 | -0.00015 | 0.00001 | 0.0751 | 0.00027 | -0.00001 | 0.00055 | 0.0545 | 0.00027 | -0.00003 | 0.00057 | 0.0803 |
| Body of corpus callosum | -0.00001 | -0.00018 | 0.00016 | 0.8943 | 0.00056 | 0.00034 | 0.00079 | <0.0001 | 0.00008 | -0.00024 | 0.00041 | 0.6075 |
| Cerebral Peduncle L | -0.00014 | -0.00022 | -0.00007 | 0.0002 | 0.00065 | 0.00038 | 0.00093 | <0.0001 | -0.00018 | -0.00048 | 0.00013 | 0.2538 |
| Cerebral Peduncle R | -0.00019 | -0.00028 | -0.00010 | <0.0001 | 0.00073 | 0.00049 | 0.00097 | <0.0001 | -0.00020 | -0.00046 | 0.00007 | 0.1420 |
| Cingulum (cingulate gyrus) L | -0.00036 | -0.00051 | -0.00021 | <0.0001 | 0.00046 | -0.00016 | 0.00107 | 0.1471 | 0.00147 | 0.00090 | 0.00203 | <0.0001 |
| Cingulum (cingulate gyrus) R | -0.00034 | -0.00046 | -0.00023 | <0.0001 | 0.00030 | -0.00033 | 0.00094 | 0.3470 | 0.00144 | 0.00086 | 0.00202 | <0.0001 |
| Cingulum (hippocampus) L | -0.00060 | -0.00078 | -0.00043 | <0.0001 | 0.00147 | 0.00080 | 0.00214 | <0.0001 | 0.00045 | -0.00022 | 0.00113 | 0.1887 |
| Cingulum (hippocampus) R | -0.00070 | -0.00086 | -0.00054 | <0.0001 | 0.00155 | 0.00100 | 0.00210 | <0.0001 | 0.00020 | -0.00037 | 0.00077 | 0.4911 |
| Corticospinal Tract L | -0.00003 | -0.00013 | 0.00006 | 0.4919 | 0.00029 | 0.00003 | 0.00055 | 0.0316 | -0.00015 | -0.00050 | 0.00019 | 0.3790 |
| Corticospinal Tract R | -0.00001 | -0.00011 | 0.00008 | 0.8072 | 0.00006 | -0.00015 | 0.00026 | 0.5906 | 0.00008 | -0.00022 | 0.00039 | 0.5890 |
| External Capsule L | -0.00030 | -0.00039 | -0.00022 | <0.0001 | 0.00035 | 0.00007 | 0.00063 | 0.0137 | 0.00044 | 0.00015 | 0.00073 | 0.0033 |
| External Capsule R | -0.00040 | -0.00049 | -0.00030 | <0.0001 | 0.00060 | 0.00030 | 0.00090 | 0.0001 | 0.00050 | 0.00019 | 0.00080 | 0.0013 |
| Fornix | 0.00032 | -0.00068 | 0.00133 | 0.5272 | 0.00302 | 0.00240 | 0.00364 | <0.0001 | -0.00172 | -0.00301 | -0.00044 | 0.0084 |
| Fornix / Stria terminalis L | -0.00049 | -0.00069 | -0.00028 | <0.0001 | 0.00141 | 0.00093 | 0.00189 | <0.0001 | 0.00029 | -0.00013 | 0.00071 | 0.1732 |
| Fornix / Stria terminalis R | -0.00069 | -0.00095 | -0.00043 | <0.0001 | 0.00201 | 0.00158 | 0.00243 | <0.0001 | 0.00001 | -0.00039 | 0.00040 | 0.9791 |
| Genu of Corpus Callosum | -0.00022 | -0.00060 | 0.00015 | 0.2466 | 0.00046 | 0.00017 | 0.00075 | 0.0021 | 0.00031 | -0.00010 | 0.00073 | 0.1327 |
| Inferior Cerebellar Peduncle L | 0.00021 | 0.00012 | 0.00030 | <0.0001 | 0.00047 | 0.00009 | 0.00085 | 0.0155 | -0.00019 | -0.00064 | 0.00026 | 0.4155 |
| Inferior Cerebellar Peduncle R | 0.00018 | 0.00010 | 0.00026 | <0.0001 | 0.00030 | -0.00008 | 0.00068 | 0.1177 | 0.00000 | -0.00044 | 0.00044 | 0.9922 |
| Medial Lemniscus L | 0.00021 | 0.00006 | 0.00037 | 0.0053 | 0.00123 | 0.00095 | 0.00150 | <0.0001 | -0.00097 | -0.00134 | -0.00061 | <0.0001 |
| Medial Lemniscus R | 0.00018 | 0.00006 | 0.00029 | 0.0031 | 0.00104 | 0.00080 | 0.00127 | <0.0001 | -0.00080 | -0.00114 | -0.00047 | <0.0001 |
| Middle Cerebellar Peduncle | 0.00004 | -0.00005 | 0.00013 | 0.3362 | 0.00055 | 0.00032 | 0.00078 | <0.0001 | -0.00014 | -0.00045 | 0.00017 | 0.3745 |
| Pontine Crossing Tract | -0.00003 | -0.00013 | 0.00007 | 0.5226 | 0.00068 | 0.00047 | 0.00089 | <0.0001 | -0.00030 | -0.00059 | -0.00002 | 0.0354 |
| Posterior Corona Radiata L | -0.00007 | -0.00028 | 0.00014 | 0.5170 | 0.00149 | 0.00104 | 0.00193 | <0.0001 | -0.00059 | -0.00113 | -0.00005 | 0.0333 |
| Posterior Corona Radiata R | -0.00003 | -0.00023 | 0.00017 | 0.7881 | 0.00158 | 0.00116 | 0.00201 | <0.0001 | -0.00080 | -0.00133 | -0.00026 | 0.0036 |
| Posterior Limb of Internal Capsule L | -0.00007 | -0.00012 | -0.00001 | 0.0155 | 0.00033 | 0.00017 | 0.00048 | <0.0001 | 0.00001 | -0.00016 | 0.00017 | 0.9268 |
| Posterior Limb of Internal Capsule R | -0.00009 | -0.00015 | -0.00003 | 0.0046 | 0.00038 | 0.00023 | 0.00053 | <0.0001 | 0.00001 | -0.00014 | 0.00017 | 0.8867 |
| Posterior Thalamic Radiation L | -0.00052 | -0.00099 | -0.00006 | 0.0260 | 0.00103 | 0.00074 | 0.00132 | <0.0001 | 0.00009 | -0.00041 | 0.00059 | 0.7325 |
| Posterior Thalamic Radiation R | -0.00018 | -0.00046 | 0.00010 | 0.2051 | 0.00096 | 0.00070 | 0.00122 | <0.0001 | -0.00022 | -0.00061 | 0.00018 | 0.2756 |
| Retrolenticular Part of Internal Capsule L | -0.00014 | -0.00026 | -0.00002 | 0.0216 | 0.00053 | 0.00031 | 0.00075 | <0.0001 | 0.00018 | -0.00010 | 0.00045 | 0.2007 |
| Retrolenticular Part of Internal Capsule R | -0.00011 | -0.00026 | 0.00003 | 0.1136 | 0.00050 | 0.00031 | 0.00069 | <0.0001 | 0.00009 | -0.00018 | 0.00035 | 0.5093 |
| Sagittal Stratum L | -0.00027 | -0.00054 | 0.00001 | 0.0545 | 0.00139 | 0.00090 | 0.00189 | <0.0001 | 0.00031 | -0.00016 | 0.00078 | 0.1992 |
| Sagittal Stratum R | -0.00018 | -0.00038 | 0.00002 | 0.0775 | 0.00101 | 0.00058 | 0.00144 | <0.0001 | 0.00040 | 0.00000 | 0.00081 | 0.0512 |
| Splenium of Corpus Callosum | -0.00016 | -0.00033 | 0.00001 | 0.0651 | 0.00041 | 0.00016 | 0.00065 | 0.0011 | 0.00030 | -0.00003 | 0.00062 | 0.0736 |
| Superior Cerebellar Peduncle L | -0.00019 | -0.00052 | 0.00013 | 0.2364 | 0.00145 | 0.00116 | 0.00174 | <0.0001 | -0.00050 | -0.00088 | -0.00011 | 0.0116 |
| Superior Cerebellar Peduncle R | -0.00027 | -0.00056 | 0.00001 | 0.0554 | 0.00143 | 0.00112 | 0.00173 | <0.0001 | -0.00044 | -0.00081 | -0.00007 | 0.0211 |
| Superior Corona Radiata L | -0.00001 | -0.00014 | 0.00011 | 0.8214 | 0.00077 | 0.00047 | 0.00107 | <0.0001 | -0.00012 | -0.00047 | 0.00023 | 0.4961 |
| Superior Corona Radiata R | 0.00000 | -0.00009 | 0.00009 | 0.9594 | 0.00064 | 0.00035 | 0.00094 | <0.0001 | -0.00022 | -0.00055 | 0.00011 | 0.1967 |
| Superior Fronto-Occipital Fasciculus L | 0.00001 | -0.00023 | 0.00025 | 0.9299 | 0.00046 | -0.00013 | 0.00105 | 0.1287 | -0.00019 | -0.00086 | 0.00048 | 0.5834 |
| Superior Fronto-Occipital Fasciculus R | 0.00002 | -0.00007 | 0.00011 | 0.7018 | 0.00041 | -0.00004 | 0.00085 | 0.0724 | -0.00031 | -0.00080 | 0.00018 | 0.2137 |
| Superior Longitudinal Fasciculus L | 0.00001 | -0.00008 | 0.00011 | 0.7917 | 0.00128 | 0.00099 | 0.00156 | <0.0001 | -0.00055 | -0.00089 | -0.00021 | 0.0017 |
| Superior Longitudinal Fasciculus R | -0.00003 | -0.00013 | 0.00008 | 0.6164 | 0.00097 | 0.00067 | 0.00128 | <0.0001 | -0.00022 | -0.00056 | 0.00012 | 0.2104 |
| Tapetum L | -0.00186 | -0.00396 | 0.00025 | 0.0836 | 0.00148 | 0.00102 | 0.00194 | <0.0001 | 0.00102 | -0.00096 | 0.00299 | 0.3131 |
| Tapetum R | -0.00138 | -0.00310 | 0.00034 | 0.1168 | 0.00095 | 0.00052 | 0.00137 | <0.0001 | 0.00083 | -0.00086 | 0.00252 | 0.3336 |
| Uncinate Fasciculus L | -0.00029 | -0.00050 | -0.00009 | 0.0051 | 0.00266 | 0.00179 | 0.00353 | <0.0001 | -0.00048 | -0.00125 | 0.00029 | 0.2222 |
| Uncinate Fasciculus R | -0.00050 | -0.00072 | -0.00029 | <0.0001 | 0.00211 | 0.00135 | 0.00288 | <0.0001 | -0.00014 | -0.00088 | 0.00060 | 0.7145 |

**Table S2:**
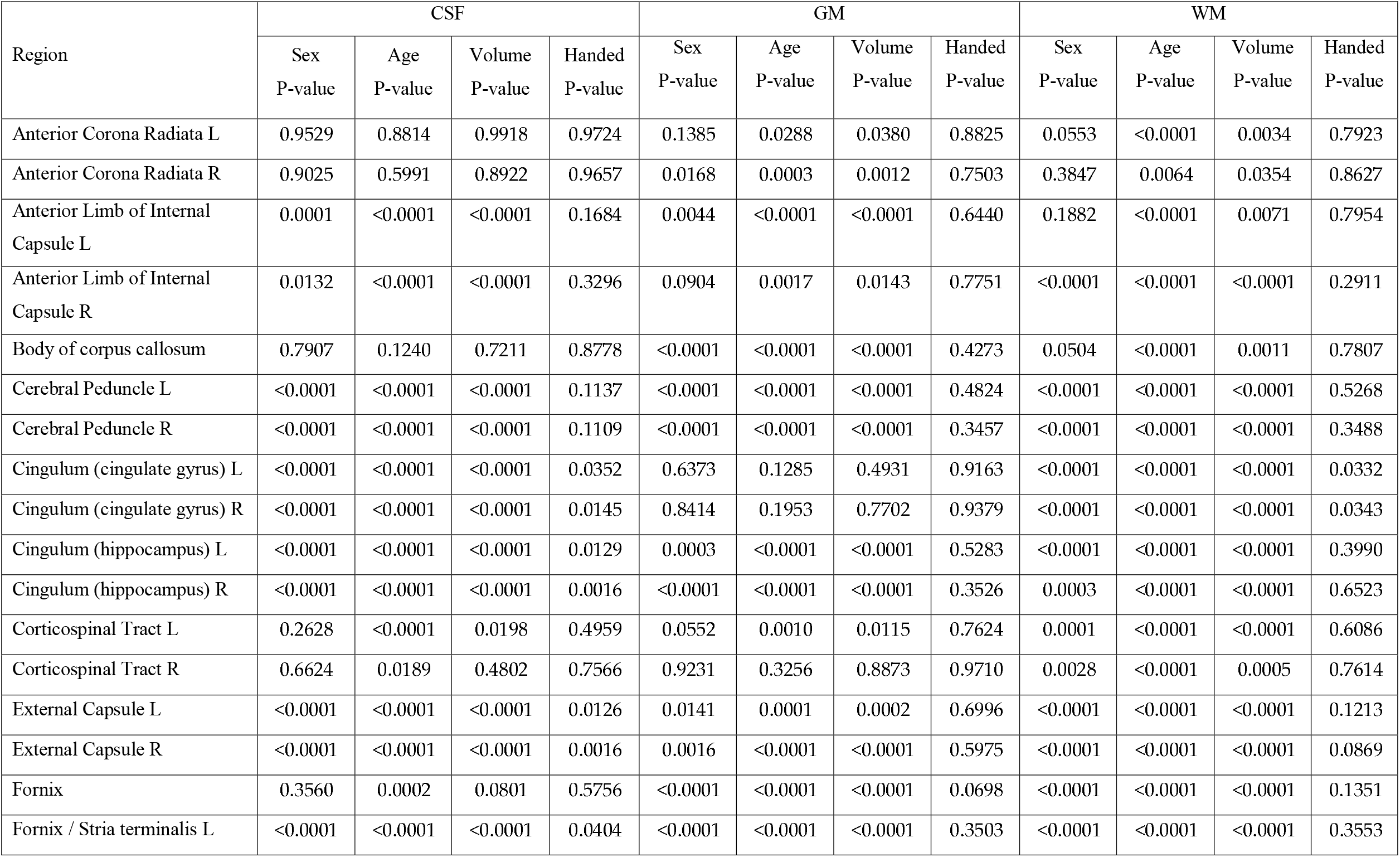
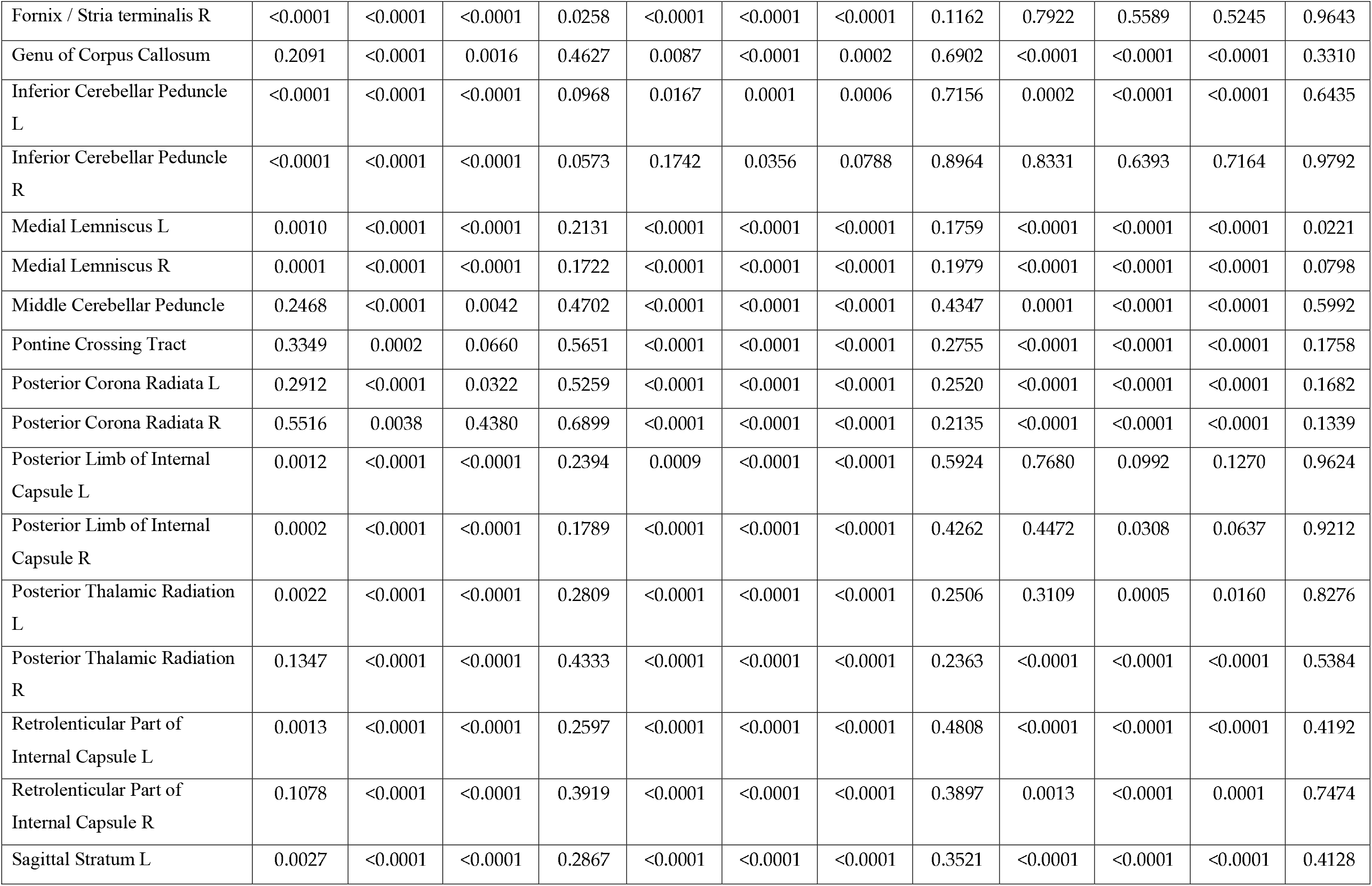
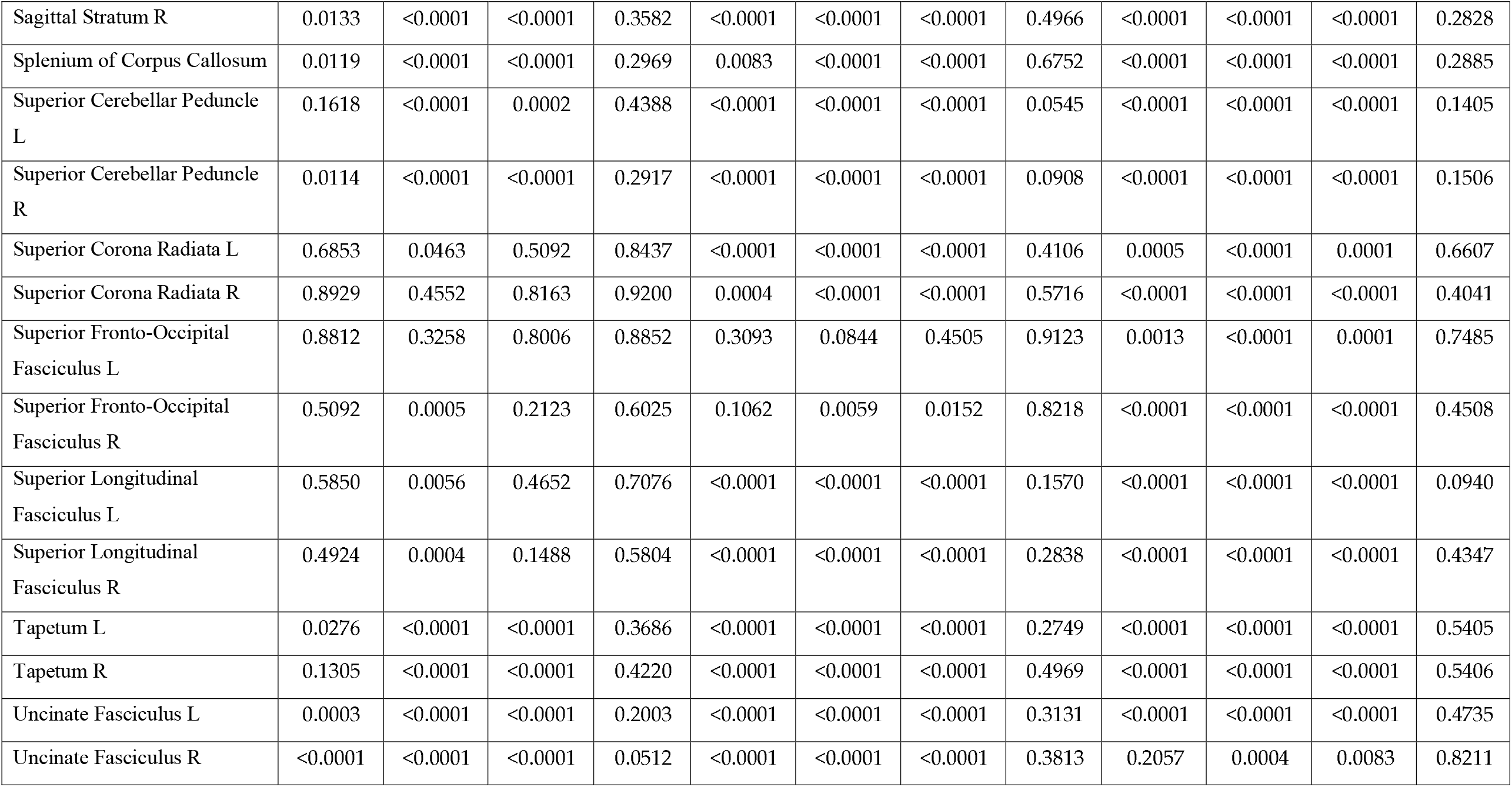
Adjusted p-values for the concomitant covariates in the models for predicting tissue signal fraction in each JHU atlas ROI.

## References

1. Lerner RM, Steinberg L. Handbook of Adolescent Psychology, Volume 1: Individual Bases of Adolescent Development. Vol 1. John Wiley & Sons; 2009.

2. Juraska JM, Willing J. Pubertal onset as a critical transition for neural development and cognition. Brain Research. 2017;1654:87–94.

3. Kessler RC, Berglund P, Demler O, Jin R, Merikangas KR, Walters EE. Lifetime prevalence and age-of-onset distributions of DSM-IV disorders in the National Comorbidity Survey Replication. Archives of general psychiatry. 2005;62(6):593–602.

4. Paus T, Keshavan M, Giedd JN. Why do many psychiatric disorders emerge during adolescence? Nature reviews neuroscience. 2008;9(12):947–957.

5. Cahill L. Why sex matters for neuroscience. Nature reviews neuroscience. 2006;7(6):477–484.

6. Nave KA. Myelination and support of axonal integrity by glia. Nature. 2010;468(7321):244–252.

7. Sisk CL, Zehr JL. Pubertal hormones organize the adolescent brain and behavior. Frontiers in neuroendocrinology. 2005;26(3-4):163–174.

8. Ahmed EI, Zehr JL, Schulz KM, Lorenz BH, DonCarlos LL, Sisk CL. Pubertal hormones modulate the addition of new cells to sexually dimorphic brain regions. Nature neuroscience. 2008;11(9):995–997.

9. Orije J, Cardon E, De Groof G, et al. In vivo online monitoring of testosterone-induced neuroplasticity in a female songbird. Hormones and behavior. 2020;118:104639.

10. Chen Z, Ye R, Goldman SA. Testosterone modulation of angiogenesis and neurogenesis in the adult songbird brain. Neuroscience. 2013;239:139–148.

11. Lebel C, Beaulieu C. Longitudinal development of human brain wiring continues from childhood into adulthood. Journal of Neuroscience. 2011;31(30):10937–10947.

12. Lebel C, Deoni S. The development of brain white matter microstructure. Neuroimage. 2018;182:207–218.

13. Peper JS, Pol HH, Crone EA, Van Honk J. Sex steroids and brain structure in pubertal boys and girls: a mini-review of neuroimaging studies. Neuroscience. 2011;191:28–37.

14. Huttenlocher PR. Synaptic density in human frontal cortex-developmental changes and effects of aging. Brain Res. 1979;163(2):195–205.

15. Natu VS, Gomez J, Barnett M, et al. Apparent thinning of human visual cortex during childhood is associated with myelination. Proceedings of the National Academy of Sciences. 2019;116(41):20750–20759.

16. Asato MR, Terwilliger R, Woo J, Luna B. White matter development in adolescence: a DTI study. Cerebral cortex. 2010;20(9):2122–2131.

17. Lebel C, Walker L, Leemans A, Phillips L, Beaulieu C. Microstructural maturation of the human brain from childhood to adulthood. Neuroimage. 2008;40(3):1044–1055.

18. Simmonds DJ, Hallquist MN, Asato M, Luna B. Developmental stages and sex differences of white matter and behavioral development through adolescence: a longitudinal diffusion tensor imaging (DTI) study. Neuroimage. 2014;92:356–368.

19. Jeurissen B, Leemans A, Tournier JD, Jones DK, Sijbers J. Investigating the prevalence of complex fiber configurations in white matter tissue with diffusion magnetic resonance imaging. Human brain mapping. 2013;34(11):2747–2766.

20. Wiegell MR, Larsson HB, Wedeen VJ. Fiber crossing in human brain depicted with diffusion tensor MR imaging. Radiology. 2000;217(3):897–903.

21. Zhang H, Schneider T, Wheeler-Kingshott CA, Alexander DC. NODDI: practical in vivo neurite orientation dispersion and density imaging of the human brain. Neuroimage. 2012;61(4):1000–1016.

22. Genc S, Malpas CB, Holland SK, Beare R, Silk TJ. Neurite density index is sensitive to age related differences in the developing brain. Neuroimage. 2017;148:373–380.

23. Mah A, Geeraert B, Lebel C. Detailing neuroanatomical development in late childhood and early adolescence using NODDI. PLoS One. 2017;12(8):e0182340.

24. Raffelt D, Dhollander T, Tournier JD, et al. Bias field correction and intensity normalisation for quantitative analysis of apparent fibre density. In: Proc. Intl. Soc. Mag. Reson. Med. Vol 25.; 2017:3541.

25. Raffelt DA, Smith RE, Ridgway GR, et al. Connectivity-based fixel enhancement: Whole-brain statistical analysis of diffusion MRI measures in the presence of crossing fibres. Neuroimage. 2015;117:40–55.

26. Genc S, Malpas CB, Gulenc A, et al. Longitudinal patterns of white matter fibre density and morphology in children are associated with age and pubertal stage. Developmental cognitive neuroscience. 2020;45:100853.

27. Silk TJ, Genc S, Anderson V, et al. Developmental brain trajectories in children with ADHD and controls: a longitudinal neuroimaging study. BMC psychiatry. 2016;16(1):1–9.

28. Petersen AC, Crockett L, Richards M, Boxer A. A self-report measure of pubertal status: Reliability, validity, and initial norms. Journal of youth and adolescence. 1988;17(2):117–133.

29. Shirtcliff EA, Dahl RE, Pollak SD. Pubertal development: correspondence between hormonal and physical development. Child development. 2009;80(2):327–337.

30. Hua K, Zhang J, Wakana S, et al. Tract probability maps in stereotaxic spaces: analyses of white matter anatomy and tract-specific quantification. Neuroimage. 2008;39(1):336–347.

31. Mori S, Wakana S, Zijl PCM van, Nagae-Poetscher LM. MRI Atlas of Human White Matter. Elsevier; 2005.

32. Wakana S, Caprihan A, Panzenboeck MM, et al. Reproducibility of quantitative tractography methods applied to cerebral white matter. Neuroimage. 2007;36(3):630–644.

33. Mito R, Dhollander T, Xia Y, et al. In vivo microstructural heterogeneity of white matter lesions in healthy elderly and Alzheimer’s disease participants using tissue compositional analysis of diffusion MRI data. NeuroImage: Clinical. 2020;28:102479.

34. Newman BT, Dhollander T, Reynier KA, Panzer MB, Druzgal TJ. Test–retest reliability and long-term stability of three-tissue constrained spherical deconvolution methods for analyzing diffusion MRI data. Magn Reson Med. 2020;84(4):2161–2173. doi:10.1002/mrm.28242

35. Iacono WG, Heath AC, Hewitt JK, et al. The utility of twins in developmental cognitive neuroscience research: How twins strengthen the ABCD research design. Developmental cognitive neuroscience. 2018;32:30–42.

36. Casey BJ, Cannonier T, Conley MI, et al. The adolescent brain cognitive development (ABCD) study: imaging acquisition across 21 sites. Developmental cognitive neuroscience. 2018;32:43–54.

37. Hagler Jr DJ, Hatton S, Cornejo MD, et al. Image processing and analysis methods for the Adolescent Brain Cognitive Development Study. Neuroimage. 2019;202:116091.

38. Veraart J, Fieremans E, Novikov DS. Diffusion MRI noise mapping using random matrix theory. Magnetic resonance in medicine. 2016;76(5):1582–1593.

39. Kellner E, Dhital B, Kiselev VG, Reisert M. Gibbs-ringing artifact removal based on local subvoxel-shifts. Magnetic resonance in medicine. 2016;76(5):1574–1581.

40. Andersson JL, Skare S, Ashburner J. How to correct susceptibility distortions in spin-echo echo-planar images: application to diffusion tensor imaging. Neuroimage. 2003;20(2):870–888.

41. Andersson JL, Graham MS, Zsoldos E, Sotiropoulos SN. Incorporating outlier detection and replacement into a non-parametric framework for movement and distortion correction of diffusion MR images. Neuroimage. 2016;141:556–572.

42. Andersson JL, Sotiropoulos SN. An integrated approach to correction for off-resonance effects and subject movement in diffusion MR imaging. Neuroimage. 2016;125:1063–1078.

43. Smith SM, Jenkinson M, Woolrich MW, et al. Advances in functional and structural MR image analysis and implementation as FSL. Neuroimage. 2004;23:S208–S219.

44. Greenspan H. Super-resolution in medical imaging. The computer journal. 2009;52(1):43–63.

45. Sotiropoulos SN, Jbabdi S, Xu J, et al. Advances in diffusion MRI acquisition and processing in the Human Connectome Project. Neuroimage. 2013;80:125–143.

46. Smith SM. Fast robust automated brain extraction. Human brain mapping. 2002;17(3):143–155.

47. Dhollander T, Raffelt D, Connelly A. Unsupervised 3-tissue response function estimation from single-shell or multi-shell diffusion MR data without a co-registered T1 image. In: ISMRM Workshop on Breaking the Barriers of Diffusion MRI. Vol 5. ISMRM; 2016.

48. Dhollander T, Connelly A. A novel iterative approach to reap the benefits of multi-tissue CSD from just single-shell (+ b= 0) diffusion MRI data. In: Proc ISMRM. Vol 24. ; 2016:3010.

49. Pinto MS, Paolella R, Billiet T, et al. Harmonization of brain diffusion MRI: Concepts and methods. Frontiers in Neuroscience. Published online 2020:396.

50. Raffelt D, Tournier JD, Fripp J, Crozier S, Connelly A, Salvado O. Symmetric diffeomorphic registration of fibre orientation distributions. NeuroImage. 2011;56(3):1171–1180.

51. Newman B, Untaroiu A, Druzgal T. A Novel Diffusion Registration Method with the NTU-DSI-122 Template to Transform Free Water Signal Fraction Maps to Stereotaxic Space.; 2020.

52. Jenkinson M, Beckmann CF, Behrens TE, Woolrich MW, Smith SM. Fsl. Neuroimage. 2012;62(2):782–790.

53. Barch DM, Albaugh MD, Avenevoli S, et al. Demographic, physical and mental health assessments in the adolescent brain and cognitive development study: Rationale and description. Developmental cognitive neuroscience. 2018;32:55–66.

54. Saragosa-Harris NM, Chaku N, MacSweeney N, et al. A practical guide for researchers and reviewers using the ABCD Study and other large longitudinal datasets. Developmental cognitive neuroscience. 2022;55:101115.

55. De Lorme KC, Schulz KM, Salas-Ramirez KY, Sisk CL. Pubertal testosterone organizes regional volume and neuronal number within the medial amygdala of adult male Syrian hamsters. Brain research. 2012;1460:33–40.

56. Zhang Z, Cerghet M, Mullins C, Williamson M, Bessert D, Skoff R. Comparison of in vivo and in vitro subcellular localization of estrogen receptors α and β in oligodendrocytes. Journal of neurochemistry. 2004;89(3):674–684.

57. Menzies L, Goddings AL, Whitaker KJ, Blakemore SJ, Viner RM. The effects of puberty on white matter development in boys. Developmental cognitive neuroscience. 2015;11:116–128.

58. Herting MM, Gautam P, Spielberg JM, Dahl RE, Sowell ER. A longitudinal study: changes in cortical thickness and surface area during pubertal maturation. PLoS One. 2015;10(3):e0119774.

59. Palmer CE, Pecheva D, Iversen JR, et al. Microstructural development from 9 to 14 years: Evidence from the ABCD Study. Developmental cognitive neuroscience. 2022;53:101044.

60. White NS, Leergaard TB, D’Arceuil H, Bjaalie JG, Dale AM. Probing tissue microstructure with restriction spectrum imaging: histological and theoretical validation. Human brain mapping. 2013;34(2):327–346.

61. Tamnes CK, Roalf DR, Goddings AL, Lebel C. Diffusion MRI of white matter microstructure development in childhood and adolescence: Methods, challenges and progress. Developmental cognitive neuroscience. 2018;33:161–175.

62. Raffelt D, Tournier JD, Rose S, et al. Apparent fibre density: a novel measure for the analysis of diffusion-weighted magnetic resonance images. Neuroimage. 2012;59(4):3976–3994.

63. Raffelt DA, Tournier JD, Smith RE, et al. Investigating white matter fibre density and morphology using fixel-based analysis. Neuroimage. 2017;144:58–73.

64. Barendse ME, Simmons JG, Smith RE, Seal ML, Whittle S. Adrenarcheal hormone-related development of white matter during late childhood. Neuroimage. 2020;223:117320.

65. Paus T. Growth of white matter in the adolescent brain: myelin or axon? Brain and cognition. 2010;72(1):26–35.

66. Willing J, Juraska JM. The timing of neuronal loss across adolescence in the medial prefrontal cortex of male and female rats. Neuroscience. 2015;301:268–275.

67. Perrin JS, Hervé PY, Leonard G, et al. Growth of white matter in the adolescent brain: role of testosterone and androgen receptor. Journal of Neuroscience. 2008;28(38):9519–9524.

